# Substantia nigra pars reticulata regulates prefrontal cortex via ventromedial thalamus

**DOI:** 10.64898/2026.01.13.699372

**Authors:** Sanne M. Casello, Adam G. Carter

## Abstract

The ventromedial thalamus (VM) innervates layer 1 (L1) of the medial prefrontal cortex (mPFC) to influence executive function and arousal. Thalamocortical (TC) cells in VM process inputs from cortex that are important for generating persistent activity in closed reciprocal loops. However, little is known about connectivity and influence of subcortical inputs to TC cells, and how they are routed through VM to circuits in the mPFC. Here we use anatomical tracing, electrophysiology, and optogenetics to investigate subcortico-thalamo-cortical circuits in mouse VM. We first characterize the morphology and physiology of TC cells in VM and determine their main subcortical input arises from substantia nigra pars reticulata (SNr). We then show how SNr inputs make strong inhibitory connections onto TC cells, which are mediated by GABA_A_ receptors and effectively suppress action potential firing. Lastly, we utilize intersectional approaches to show how SNr inputs are channeled via TC cells in VM to engage specific inhibitory networks in L1 of mPFC. Together, our results indicate how subcortical inputs engage higher-order thalamus to influence the frontal cortex, highlighting differences from equivalent circuits in motor systems.

**SIGNIFICANCE STATEMENT:** Here we use anatomy, slice physiology, and optogenetics to examine how subcortical inputs are routed through the ventral medial thalamus (VM) to the medial prefrontal cortex (mPFC). We first find that the main subcortical input to thalamocortical (TC) cells arises from substantia nigra pars reticulata (SNr), rather than cerebellum or internal globus pallidus. We then use electrophysiology and optogenetics to show that SNr input is inhibitory, mediated by GABA_A_ receptors, and sufficient to silence TC cell firing. Lastly, we demonstrate that SNr inputs are channeled via TC cells in VM to engage NDNF+ interneurons in layer 1 (L1) of the mPFC. Together, our findings illustrate how subcortical inputs can signal through higher-order thalamus to influence frontal cortical circuits.

## INTRODUCTION

Higher-order thalamus communicates with the medial prefrontal cortex (mPFC) to guide cognitive and motivated behaviors (Euston *et al*., 2012; Halassa & Sherman, 2019). Disruption of these connections is closely linked to neuropsychiatric disorders, including schizophrenia, depression, and anxiety (Euston *et al*., 2012; Huang *et al*., 2019; Xu *et al*., 2019; Klein-Flugge *et al*., 2022). We previously found that the ventromedial (VM) thalamus makes strong, reciprocal connections with the mPFC (Collins *et al*., 2018). However, the ability of other inputs from subcortical brain regions to engage VM, and thus influence the mPFC, remains largely unexplored. Investigating these connections is important, in order to better understand how dysregulation of these circuits leads to disruption of complex behaviors associated with many neuropsychiatric disorders.

The VM is classically defined as “matrix” thalamus (Jones, 1998; 2001), with thalamocortical (TC) cells that diffusely project to superficial layers of the cortex (Kuramoto *et al*., 2015; Halassa & Sherman, 2019; Shepherd & Yamawaki, 2021). While TC cells in “core” thalamus can effectively drive cortical circuits (Kuramoto *et al*., 2017), TC cells in matrix thalamus have a more regulatory influence (Glenn & Steriade, 1982; Chen *et al*., 2015). Motor cortex is a major cortical target of VM (Guo *et al*., 2018), where these connections influence motor planning and execution (Komiyama *et al*., 2010; Chen *et al*., 2017; Takahashi *et al*., 2021; Wang *et al*., 2021). mPFC also receives robust connections from VM (Cruikshank *et al*., 2012; Sieveritz *et al*., 2019; Anastasiades *et al*., 2021), which are involved in arousal (Honjoh *et al*., 2018; Iidaka, 2021). However, while connections between VM and motor cortex are well explored (Hooks *et al*., 2013; Guo *et al*., 2017), equivalent circuits linking VM and mPFC have not been extensively described.

In addition to receiving inputs from cortex, VM and other higher-order thalamic nuclei receive and process a variety of subcortical inputs (Aoki *et al*., 2019). The motor cortex-projecting parts of VM receive inputs from the basal ganglia and cerebellum (Gao *et al*., 2018; Inagaki *et al*., 2022; Alonso-Martinez *et al*., 2023). The substantia nigra pars reticulata (SNr) sends strong, GABAergic projections to VM (Beckstead *et al*., 1979; Deniau *et al*., 1994; Kase *et al*., 2015; Lobb & Jaeger, 2015; Lee *et al*., 2020), which shape thalamic output to motor cortex to dictate movement initiation (Catanese & Jaeger, 2021). In contrast, the cerebellum (CbN) sends glutamatergic projections across the lateral to medial axis of ventral thalamus, with strong innervation of the ventral lateral (VAL) thalamus (Gornati *et al*., 2018), and activation of CbN has been shown to effectively control thalamo-cortical network activity (Proville *et al*., 2014; Kros *et al*., 2015; Gornati *et al*., 2018). However, the extent and properties of equivalent long-range connections from different subcortical brain regions to mPFC-projecting TC cells in VM remain relatively unexplored.

Ultimately, TC cells in VM densely innervate the outer edge of layer 1 (L1), the most superficial layer of mPFC (Cruikshank *et al*., 2012; Anastasiades *et al*., 2021). L1 contains the dendrites of pyramidal cells, as well as GABAergic interneurons that express neuron-derived neurotrophic factor (NDNF) (Rudy *et al*., 2011; Schuman *et al*., 2019; Anastasiades *et al*., 2021). L1 NDNF+ cells display late spiking, high input resistance, expansive horizontal dendritic tuffs, and prolific axonal arborizations (Hartung *et al*., 2024). In many cortices, NDNF+ cells play a key role in shaping cortical dynamics and thus arousal and learning (Cohen-Kashi Malina *et al*., 2021; Ibrahim *et al*., 2021). In the mPFC, VM inputs preferentially target and trigger action potential firing at NDNF+ cells over other interneurons, and NDNF+ cells in turn target PV+ and SOM+ interneurons in other layers to engage disinhibitory networks (Huang *et al*., 2024). In principle, subcortical inputs may access these inhibitory circuits in L1 of the mPFC via TC cells in VM, but establishing the presence of these kinds of polysynaptic neural circuits has proven challenging.

Here, we combine anatomical tracing, electrophysiology, and optogenetics to dissect how subcortical inputs are routed via VM to the mPFC. We first characterize the distribution, morphology, and physiology of mPFC-projecting TC cells in VM. We then establish that SNr is the main subcortical input to these cells, with minimal input from the CbN. We next use anatomy and physiology to characterize SNr inputs, including their sign, dynamics, and functional impact. Lastly, we examine how SNr connections are routed via VM to influence L1 of the mPFC, showing how they can ultimately engage NDNF+ interneurons. Together, our findings indicate how subcortical inputs can impact TC cells in VM to engage neural circuits in the mPFC, with implications for cognitive and motivational behaviors, as well as neuropsychiatric disorders.

## MATERIALS & METHODS

### Animals

Experiments used P30 – P84 wild-type (WT) and transgenic mice of either sex bred on a C57BL/6J background (all breeders from Jackson Labs or Mutant Mouse Resource and Research Centers). No animals had been involved in previous procedures. Animals were group-housed with same-sex littermates in a dedicated animal care facility and were on a 12-h light/dark cycle at 18 – 23°C. Food and water were available *ad libitum*. Transgenic mice included: NDNF-Cre (JAX 030757; Tasic et al., 2016), Ai14 (Cre-dependent tdTomato; JAX 007914; Madisen *et al*., 2010), GAD2-cre (JAX 028867; Taniguchi *et al*., 2011), Vglut2-cre (JAX 028863; Vong *et al*., 2011), and Syt6-Cre (MMRRC #037416; Gong *et al*., 2003). Homozygote male breeders were paired with female wild-type breeders to yield heterozygote offspring used for experiments. All experimental procedures were approved by the University Animal Welfare Committee of New York University.

### Viruses

Adeno-associated viruses (AAVs) and rabies viruses (RVs) used in this study included: AAV1-hSyn-hChR2(H134R)-eYFP (UPenn # AV-26973P), AAV1-hSyn-Cre (Addgene # 105553), AAV1-Ef1a-Flpo (Addgene # 55637), AAV1-hSyn-Chrimson-tdTomato (Addgene # 59171), AAV1-EF1a-DIO-eYFP (Addgene # 27056), AAV1-CAMKII-ChR2(H134R)-mCherry (Addgene # 26975) AAV1-CAG-FLEX-tdTomato (Addgene # 28306), AAV1-Syn-Chronos-GFP (UPenn # AV-1-PV3446), AAV1-CAG-FLEXFRT-ChR2-mCherry (Addgene # 75470), AAV1-hSyn-FLEX-mGFP-2A-Synaptophysin-mRuby (Addgene # 71760), CVS-N2c(dG)-H2B-tdTomato (UNC Neurotools #241118-2), CVS-N2c(dG)-H2B-GFP (UNC Neurotools #241118-2), AAVrg-CAG-H2B-EGFP (UNC Neurotools # NT23576), AAV2-retro-Syn-Cre (UNC Neurotools # NT231051.1), AAV1-Syn-DIO-TVA66T-dTomato-CVS-N2cG (UNC Neurotools # NT23521), EnvA-CVS-N2c(dG)-H2B-EGFP (UNC Neurotools # NT2357).

### Stereotaxic injections

Mice aged 4 – 6 weeks were deeply anesthetized with 3 – 4% isoflurane in an induction chamber, maintained at 1 – 3% isoflurane, and head-fixed in a stereotaxic frame (Kopf Instruments). A small craniotomy was made over the injection site, using these coordinates relative to Bregma: mPFC A/P = 2.1 mm, M/L = 0.35, D/V = -2.05 & 2.25. VM (30° angle) A/P = -1.2 mm, M/L = 2.9 mm D/V = -3.6 mm. SNr A/P = -3.32 mm, M/L = -1.2 & -1.5mm, D.V = -4.5 mm & -4.7 mm. CbN A/P = -6.0 mm, M/L = -1.5 mm, D/V = -3.6 & -3.8 mm. Borosilicate pipettes with 5 to 10 µm diameter tips were back-filled with 100 – 275 nL of virus and pressure-injected using a Nanoject III (Drummond) every 45 s. The pipette was left in place for an additional 10 min, allowing time to diffuse away from the pipette tip, before being slowly retracted from the brain. For both retrograde and viral labeling, animals were housed for 2 – 3 weeks before slicing. For slice physiology experiments, animals were housed for 3 – 4 weeks before slicing. For trans-neuronal anterograde tracing and monosynaptic rabies tracing experiments, animals were housed for 6 – 10 weeks before slicing.

### Cell-type specific rabies virus tracing

For monosynaptic rabies virus tracing, AAVrg-Cre virus mixed with cholera toxin subunit B conjugated to Alexa 647 (CTB-647) (4:1 dilution, 125 nL total) was injected into a single hemisphere of the mPFC in WT mice. Concurrently, AAV-DIO-TVA-N2cG-tdTomato helper virus (274.5 nL) was injected into VM. After allowing 5 weeks for expression of helper viruses, EnvA-RV-ΔG-GFP rabies virus (274.5 nL) was injected at the same location in VM. After an additional 8 days to allow for monosynaptic retrograde labeling, mice were perfused, and slices prepared for fluorescent microscopy, as described below.

### Histology and fluorescence microscopy

Mice were anesthetized with 3 – 4% isoflurane in an induction chamber and maintained at 1 – 3% isoflurane. They were then perfused intracardially with 0.01 M phosphate-buffered saline (PBS) followed by 4% paraformaldehyde (PFA) in 0.01 M PBS. Brains were fixed in 4% PFA in 0.01 M PBS overnight at 4°C. Slices were prepared at a thickness of 70 μm (Leica VT 1000S vibratome) and directly mounted onto gel-coated glass slides. For biocytin-filled cells, slices were also stained with streptavidin conjugated Alexa 647. Slices were cover-slipped using VectaShield with DAPI (Vector Labs). Fluorescent images were taken on an Olympus VS120 microscope, using either a 10 x 0.25 NA objective (Olympus) or a Leica TCS SP8 confocal microscope using a 10x 0.4 NA, 20x 0.75 NA, or 63x 1.4 NA oil-immersion objective (Leica). Image processing involved adjusting brightness and contrast using ImageJ (NIH). Cell counting of nuclear-localized fluorescent labeling was performed using semi-automated cell detection software NeuroInfo (MBF Bioscience). Cell counting of cytoplasmic cellular labeling was performed manually using ImageJ (NIH). Axon fluorescence analysis and heatmaps were generated using ImageJ (NIH) and images were aligned to the Allen Common Coordinate Framework (Wang *et al*., 2020b).

### Slice preparation

Mice aged 6 – 8 weeks were anesthetized with 3 – 4% isoflurane in an induction chamber, maintained at 1 – 3% isoflurane, and perfused intracardially with an ice-cold external solution containing the following (in mM): 65 sucrose, 76 NaCl, 25 NaHCO_3_, 1.4 NaH_2_PO_4_, 25 glucose, 2.5 KCl, 7 MgCl_2_, 0.4 Na-ascorbate, and 2 Na-pyruvate (295–305 mOsm) and bubbled with 95% O_2_/5% CO_2_. Coronal slices (300 μm thick) were cut on a VS1200 vibratome (Leica) in ice -cold external solution and transferred to ACSF containing (in mM): 120 NaCl, 25 NaHCO_3_, 1.4 NaH_2_PO_4_, 21 glucose, 2.5 KCl, 2 CaCl_2_, 1 MgCl_2_, 0.4 Na-ascorbate, and 2 Na-pyruvate (295–305 mOsm), bubbled with 95% O_2_/5% CO_2_. Slices were kept for 30 min at 35°C and recovered for 30 min at room temperature before starting recordings. All recordings were conducted at 30 – 32°C.

### Electrophysiology

Whole-cell recordings were obtained from thalamocortical (TC) cells in VM thalamus or L1 neuron derived neurotrophic factor (NDNF+) interneurons in mPFC. TC cells were identified by retrograde labeling from mPFC, primarily using AAVrg-H2B-tdTomato as a marker. Cells were identified by infrared-differential interference contrast or fluorescence, as previously described (Chalifoux & Carter, 2010). Borosilicate pipettes (3 – 5 MΩ) were filled with one of two internal solutions. For current-clamp recordings (in mM): 135 K-gluconate, 7 KCl, 10 HEPES, 10 Na-phosphocreatine, 4 Mg_2_-ATP, and 0.4 Na-GTP, 290–295 mOsm, pH 7.3, with KOH. For voltage-clamp recordings (in mM): 135 Cs-gluconate, 10 HEPES, 10 Na-phosphocreatine, 4 Mg_2_-ATP, and 0.4 Na-GTP, 0.5 EGTA, 10 TEA-chloride, and 2 QX314, 290–295 mOsm, pH 7.3, with CsOH. In some experiments studying cellular morphology, 5% biocytin was also included in the recording internal solution. After allowing biocytin to diffuse through the recorded cell for at least 30 min, slices were fixed with 4% PFA before staining with streptavidin conjugated to Alexa 647 (Invitrogen). Electrophysiology recordings were made with a Multiclamp 700B amplifier (Axon Instruments), filtered at 4 kHz for current-clamp, and 2 kHz for voltage-clamp, and sampled at 10 kHz. The initial series resistance was <20 MΩ, and recordings were ended if series resistance rose above 25 MΩ. In many experiments, 10 μM CPP was used to block NMDA receptors (Tocris Bioscience).

### Optogenetics

Channelrhodopsin-2 (ChR2) was expressed in presynaptic cells and activated with a 2 ms light pulse from a blue LED (473 nm) (Thorlabs). For wide-field illumination, light was delivered via a submerged 10x 0.3 NA objective (Olympus) centered on the recorded cell, as previously described (Collins *et al*., 2018; Anastasiades *et al*., 2021; Kamalova *et al*., 2024). LED power was routinely calibrated at the back aperture of the objective, and adjusted to obtain reliable responses, with typical values ranging from 1 to 10 mW. In some cases, stimulus trains were used, typically consisting of 5 pulses of 2 ms light delivered every 100 ms (10 Hz).

### Data analysis

Electrophysiology data was acquired using National Instruments boards and custom software written in MATLAB (MathWorks) (Pologruto *et al*., 2003). Off-line analysis was performed using custom software written in Igor Pro (WaveMetrics). Input resistance was calculated from the steady-state voltage during a -50 pA, 500 ms current step. For voltage-clamp experiments with train stimulation, each postsynaptic current (PSC) amplitude was measured as the average value in a 1 ms window around the peak, minus the average 2 ms baseline value before each stimulation. For current-clamp experiments, the number of action potentials was counted in 500 ms bins before, during, and following LED delivery.

For all anatomy quantification, coronal sections were aligned to the Allen Common Coordinate Framework (CCF) (Wang *et al*., 2020b) with NeuroInfo (MBF Bioscience) in a semi-automated way. For co-localization analysis in Ai14 mice, cell counting was performed in FIJI ImageJ on an aligned, multi-color image of retrogradely labeled neurons and trans-neuronal labeled neurons. Labeled cell bodies were manually counted in VM. For anterograde tracing, axon fluorescence analysis was manually performed in FIJI ImageJ by measuring the pixel intensity on an aligned, multi-color image of axon output in VM, ventrolateral thalamus (VAL), mPFC, anterolateral motor cortex (ALM), or insular cortex (aIC). Baseline subtraction to control for background autofluorescence on pixel intensity value of the brain region of interest was done to lowest, non-zero fluorescence value in the full coronal section. For each experimental condition, three samples were analyzed. For retrograde tracing and TRIO rabies experiments, cell counting was performed using NeuroInfo (MBF Bioscience) in a semi-automated way for 4 – 5 samples.

Most summary data are reported in the text and figures as arithmetic mean ± SEM. Comparisons between data recorded in pairs were performed using non-parametric Wilcoxon test. For unpaired comparisons of more than two groups, Kruskal-Wallis tests with Dunn’s multiple comparisons tests were performed. For paired comparisons of more than two groups, Friedman tests with Dunn’s multiple comparisons tests were performed. Two-tailed p values < 0.05 (*) were considered significant.

## RESULTS

### Characterizing mPFC-projecting TC cells in VM thalamus

We used anatomy and physiology to define the location and properties of mPFC-projecting TC cells in VM thalamus. We first co-injected into the mPFC a retrogradely transported glycoprotein-deleted rabies virus that expresses histone-tagged green fluorescent protein (RV(ΔG)-GFP), along with an anterogradely transported, Cre-dependent adeno-associated virus (AAV) that expresses tdTomato fluorophore (AAV-FLEX-tdTomato) in Syt6-Cre mice **(Fig. 1A)**. After waiting for expression and transport, we imaged green cells and red axons in serial coronal sections through the thalamus, which were aligned to the common-coordinate frame (CCF) (Wang *et al*., 2020a). We found close overlap of TC cells and mPFC axons in VM thalamus (Collins *et al*., 2018), with colocalization mostly along the ventral border near the midline (**Fig. 1B)**. Aligning and registering coronal sections confirmed that TC cells are found across the anterior-posterior (A-P) axis, with a bias towards anterior VM, and prominence in ventral locations (n = 5 mice) **(Fig. 1C).**

**Figure 1.**
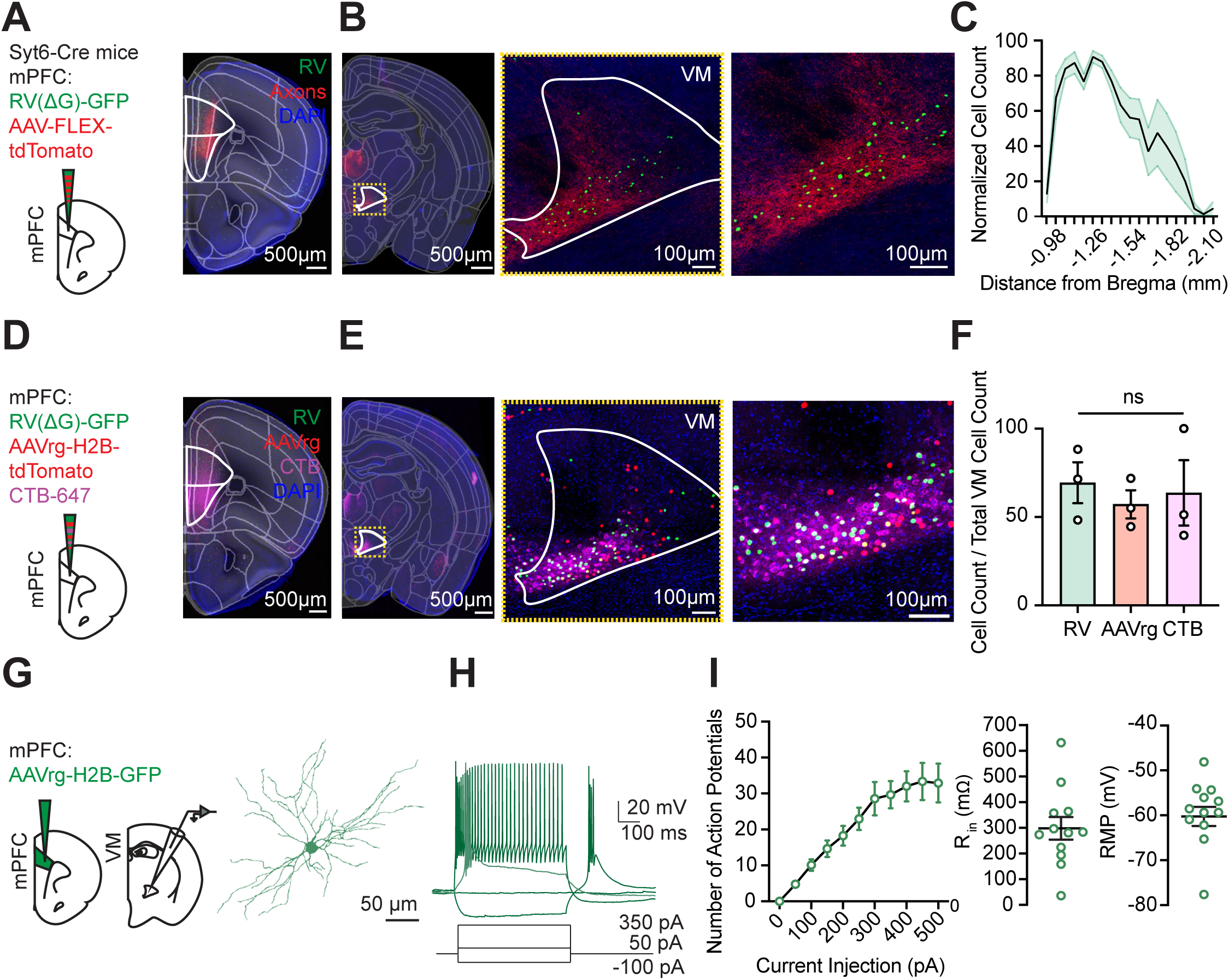
Characterization of mPFC-projecting TC Cells in VM. (**A**) *Left,* Schematic of dual retrograde and anterograde viral tracing strategy, with RV(ΔG)-H2B-GFP and AAV-FLEX-tdTomato injected into the mPFC of Syt6-Cre mice. *Right,* Representative, CCF-aligned coronal mPFC section, showing injection site with GFP+ cells (green) and tdTomato+ cells (red). mPFC is outlined in white and partitioned into prelimbic (PrL, dorsal) and infralimbic (IL, ventral) regions. Scale bar = 500 µm. **(B)** *Left,* Representative, CCF-aligned, thalamic coronal section, showing colocalization of GFP+ cells and tdTomato+ axons in VM. VM is outlined in white. Scale bar = 500 μm. *Middle,* Confocal image of magnified yellow box outline of VM. *Right,* Further magnification. Scale bar = 100 μm. **(C)** Distribution of GFP+ cells in VM across the anterior-posterior (A-P) axis. Normalized within animal to total cell count. A-P coordinate reported as millimeters from Bregma. n = 5 mice. **(D)** *Left,* Schematic of three retrograde tracing strategies, with RV(ΔG)-H2B-GFP, AAVrg-H2B-tdTomato, and CTB-647 injected into the mPFC. *Right,* Representative image of mPFC injection site, with GFP+ cells (green), tdTomato+ cells (red), and CTB-647+ cells (magenta). mPFC is outlined in white and partitioned into prelimbic (PrL, dorsal) and infralimbic (IL, ventral) regions. Scale bar = 500 μm. **(E)** *Left,* Representative, CCF-aligned, thalamic coronal section, showing GFP+, tdTomato+, and CTB-647+ cells in VM. Scale bar = 500 μm. *Middle,* Confocal image of magnified yellow box outline of VM. *Right,* Further magnification. Scale bar = 100 μm. **(F)** Relative cell counts for the three approaches, normalized within animal to total cell count in VM. Friedman test: Q = 0.7, p = 0.9. n = 3 mice. **(G)** *Left,* Schematic of AAVrg-H2B-GFP injections in mPFC for whole-cell recordings in VM. *Right,* Representative biocytin fill of GFP+ TC cell in VM. Scale bar = 50 μm. **(H)** Representative action potential firing of VM TC cell in response to current injection of 50 pA (light green trace), 350 pA, and -100 pA (dark green traces) at -65 mV. **(I)** *Left,* Summary input-output curve, showing number of action potentials fired in response to increasing injections of current. *Middle,* Summary of input resistance. *Right,* Summary of resting membrane potential. n = 11 cells, 5 mice. Values are mean ± SEM.

Although rabies virus is useful for anatomically defining the locations of presynaptic cells, electrophysiological characterization of neurons requires an alternative method. Therefore, we next compared three different ways of retrograde labeling of TC cells, including RV(ΔG)-GFP, a retrogradely transported AAV that expressed histone-tagged red fluorescent protein (AAVrg-H2B-tdTomato), and cholera toxin B tagged with Alexa fluorophore 647 (CTB-647) **(Fig. 1D).** We found that triple-labeled, CCF-registered, thalamic coronal sections showed strong labeling within VM, with similar topographies **(Fig. 1E).** Quantification of each retrograde labeling method showed a similar degree of labeling (RV = 69.4 ± 11.6 cells, AAVrg = 57.2 ± 7.9 cells, CTB = 63.7 ± 18.5 cells; Friedman test: Q = 0.7, p = 0.9; n = 3 mice) **(Fig. 1F)**, and therefore we used these methods interchangeably in our subsequent future experiments.

Having established the location of mPFC-projecting TC cells, we next assessed their intrinsic anatomy and physiology with whole-cell electrophysiology. We performed current-clamp recordings from fluorescently labeled cells in acute coronal slices while filling with biocytin for cell reconstructions (n = 11 cells, 5 mice). We found TC cells have radial dendrites **(Fig. 1G)**, which were consistent with previous studies (Varga *et al*., 2002). We also observed characteristic firing and intrinsic properties (R_in_ = 298.1 ± 43.6 MΩ, V_rest_ = -60.0 ± 2.1 mV) **(Fig. 1H-I)**, which were also consistent with prior studies (Anastasiades *et al*., 2021). Together, these findings better define the anatomical and intrinsic properties of TC cells in VM, allowing us to study their circuitry.

### Identifying putative subcortical inputs to PFC-projecting TC cells

Higher-order thalamus lacks direct input from the periphery but can receive excitatory and inhibitory afferents from subcortical sources. Previous studies on motor thalamus identified the substantia nigra pars reticulata (SNr), cerebellar nuclei (CbN), and internal globus pallidus (GPi) as major inputs to motor cortex-projecting TC cells (Catanese & Jaeger, 2021; Takahashi *et al*., 2021; Alonso-Martinez *et al*., 2023; Koster & Sherman, 2024). Therefore, we next tested whether SNr, CbN, and GPi can also provide substantial inputs to mPFC-projecting TC cells in VM.

To map the brain-wide innervation of mPFC-projecting VM cells, we used TRIO (Tracing the Relationship of Inputs and Outputs) mapping (Schwarz *et al*., 2015). This rabies-based tracing strategy enables identification of presynaptic cells to mPFC-projecting cells in VM. First, we injected a retrogradely transported AAV into the mPFC to express Cre at TC cells in VM (AAVrg-Cre). Concurrently, we injected Cre-dependent helper AAV into VM to express the TVA receptor and G glycoprotein (AAV-DIO-TVA-N2cG-tdTomato). After allowing for expression, we injected serotyped rabies virus (RV) into VM, which infects TVA-expressing TC cells and labels their presynaptic inputs (EnvA-RV-ΔG-GFP) **(Fig. 2A)**. We then prepared coronal slices, imaged fluorescence, aligned to the CCF, and used automated software to count cells. Starter cells that express both helper AAV and serotyped rabies virus were localized to ventral VM (**Fig. 2B**), and monosynaptic input cells were found across the anterior-posterior axis of the brain (n = 4 mice) **(Fig. 2C).** While most inputs to mPFC-projecting TC cells were cortical, we found that subcortical inputs made up a large fraction of the total (cortex = 66.3 ± 5.8%, subcortex = 33.7 ± 5.8%) **(Fig. 2D).** Consistent with prior studies on reciprocal thalamocortical loops, the dominant inputs to mPFC-projecting VM neurons were from the medial prefrontal cortex (mPFC = 27.3 ± 2.0%), which includes prelimbic (PrL = 10.6 ± 0.5%), infralimbic cortex (ILA = 1.7 ± 0.4%), and anterior cingulate (ACC = 15.0 ± 3.3%). Additionally, inputs from orbital (ORB = 21.7 ± 2.3%), motor cortex (MO = 14.5 ± 4.7%), and insular cortex (aIC = 6.6 ± 0.6%) were observed **(Fig. S2E-G).** The main subcortical input was SNr (SNr = 5.3 ± 1.6%), with presynaptic cells spanning the anterior-posterior axis, which mirrored that of generalized monosynaptic rabies input to VM in separate experiments (n = 4 mice) **(Fig. 2E & S2A-D).** In contrast, the cerebellar nuclei (interposed (IP) =0.3 ± 0.2%, dentate nucleus (DN) = 0.2 ± 0.1%, and fastigial (FN) = 0.2 ± 0.1%) and internal globus pallidus (GPi = 0.1 ± 0.1%) were not major inputs to mPFC-projecting TC cells (Friedman test: Q = 11.4, p = 0.007; Dunn’s multiple comparisons test: p = 0.03 for SNr vs. FN and SNr vs. GPi; n = 4 mice) **(Fig. 2F-H)**. These anatomy findings indicate that the greatest subcortical input to mPFC-projecting neurons in VM is SNr, which may signal via VM to mPFC.

**Figure 2.**
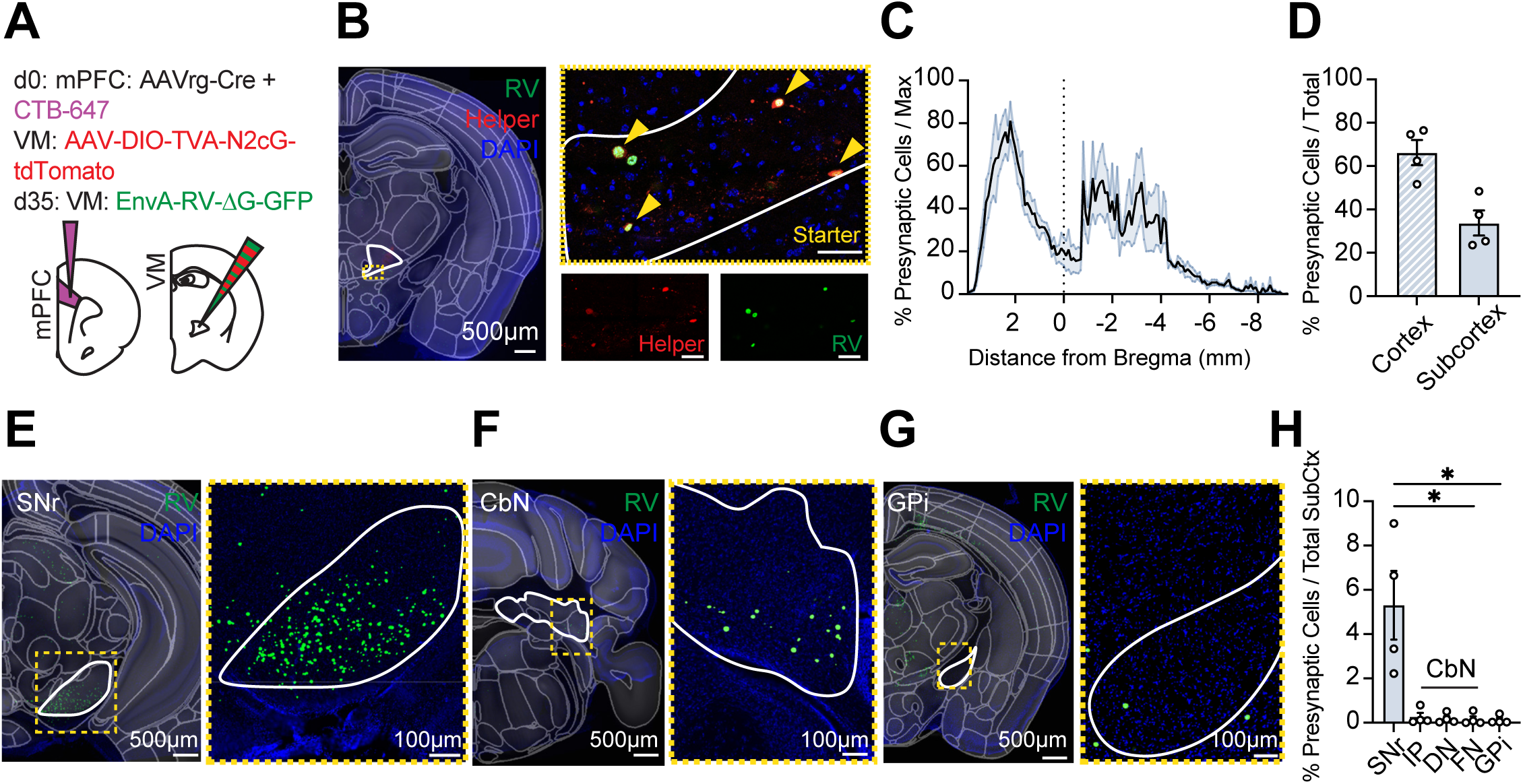
Identifying subcortical Inputs to mPFC-projecting TC cells in VM. (**A**) Schematic of monosynaptic rabies tracing, including injections of AAVrg-Cre (co-injected with a small volume of CTB-647 to mark injection location) in mPFC, helper AAV in VM, and EnvA-serotyped RV in VM to label presynaptic input cells. **(B)** *Left,* Representative, CCF-aligned, thalamic coronal section, showing tdTomato+ (red) and GFP+ (green) cells. VM is outlined in white. Scale bar = 500 μm. *Right,* Confocal images of magnified yellow box outline of VM. Split channels of tdTomato and GFP. Starter cells (yellow) are indicated by yellow arrows. Scale bar = 100 μm. **(C)** Summary distribution of GFP+ presynaptic cells along the anterior-posterior axis, reported as percentage of presynaptic cells normalized to total cell count in each animal. n = 4 mice. **(D)** Summary of GFP+ presynaptic cells in cortex and subcortex, reported as percentage of presynaptic cell count normalized to total cell count in each animal. Data points from individual mice are shown as colored circles. n = 4 mice. **(E)** *Left,* Representative, CCF-aligned, images of GFP+ presynaptic cells in SNr, the top subcortical input to TC cells in VM. SNr is outlined in white. Scale bar = 500 μm. *Right,* Cropped image of magnified yellow box outline of SNr. Scale bar = 100 μm. **(F)** Similar to (E) for CbN. **(G)** Similar to (E) for GPi. **(H)** Summary of subcortical inputs to TC cells in VM, reported as percentage of presynaptic cell count normalized reported as percentage of total cells in subcortical structures. SNr, substantia nigra pars reticulata; CbN, Cerebellar Nuclei including IP (interpositus nucleus), DN (dentate nucleus), and FN (fastigial nucleus); GPi, Internal globus pallidus. Friedman test: Q = 11.4, p = 0.007; Dunn’s multiple comparisons test: p = 0.03 for SNr vs. FN and SNr vs. GPi. n = 4 mice. Values are mean ± SEM. * = p < 0.05.

### Anatomical tracing of SNr inputs to mPFC-projecting TC cells in VM

We next used complementary anatomical strategies to assess whether SNr contacts mPFC-projecting TC cells in VM. First, we injected AAV1-Cre into the SNr of Ai14 reporter mice, in which expression of Cre recombinase drives expression of tdTomato **(Fig. 3A-B & S3A)**. At high titer, AAV1 jumps anterogradely one synapse, leading to viral expression in the postsynaptic cell (Zingg *et al*., 2017). Concurrently, we injected AAVrg-H2B-GFP into the mPFC to express histone-tagged green fluorescent protein at TC cells in the VM **(Fig. 3A-B & S3A)**. After allowing for expression and transport, we surveyed and aligned coronal slices across thalamus to examine the colocalization of red, tdTomato+ (SNr-contacted) cells and green, GFP+ (mPFC-projecting) cells **(Fig. 3C)**. We observed pronounced colocalization in VM thalamus, but not in other thalamic nuclei that project to mPFC, including mediodorsal (MD) or central medial (CM) (VM = 23.3 ± 2.6%, MD = 2.5 ± 0.4%, CM = 1.5 ± 0.4%; Friedman test: Q = 6, p = 0.03; Dunn’s multiple comparisons test: p = 0.04 for VM vs. CM; n = 3 mice) **(Fig. 3D & S3B)**.

**Figure 3:**
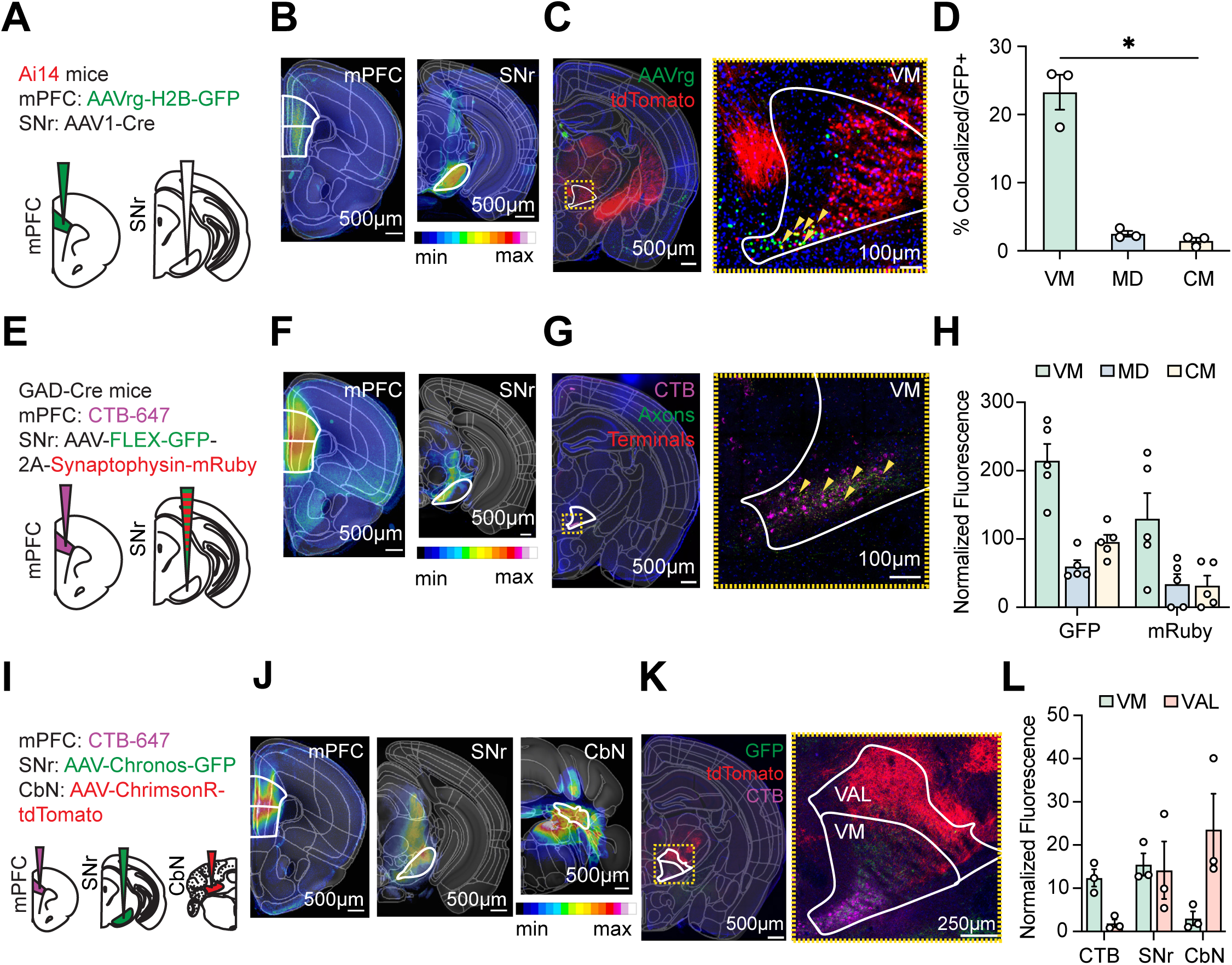
Anterograde anatomy of subcortical inputs to VM. (**A**) Schematic of AAVrg-H2B-GFP injection in the mPFC and AAV1-Cre injection in the SNr of Ai14 transgenic reporter mice, where Cre cells robustly express tdTomato. **(B)** Representative, CCF-aligned, heatmaps of mPFC and SNr injection sites with respective regions of interest outlined in white. Scale bar = 500 μm. Heatmap key shows fluorescence value. **(C)** *Left,* Representative, CCF-aligned, thalamic coronal section, showing retrogradely infected GFP+ TC cells (green) and trans-neuronally labeled tdTomato+ cells (red) in VM. VM is outlined in white. Scale bar = 500 μm. *Right,* Confocal image of magnified yellow box outline of VM, where colocalized GFP+ and tdTomato+ cells are identified with yellow arrows. Scale bar = 100 μm. **(D)** Summary of colocalized cells as a percentage of total GFP+ cells for VM, MD, and CM. Friedman test: Q = 6, p = 0.03; Dunn’s multiple comparisons test: p = 0.04 for VM vs. CM. n = 3 mice. **(E)** Schematic of CTB-647 injection into mPFC and AAV-FLEX-GFP-2A-Synaptophysin-mRuby injection into SNr of GAD-Cre mice. **(F)** Representative, CCF-aligned, heatmaps of mPFC and SNr injection sites with respective regions of interest outlined in white. Scale bar = 500 μm. Heatmap key shows fluorescence value. **(G)** *Left,* Representative, CCF-aligned, thalamic coronal section, showing retrogradely infected TC cells (magenta), GFP+ SNr axons (green), and mRuby+ SNr terminals (red). VM is outlined in white. Scale bar = 500 μm. *Right,* Confocal image of magnified yellow box outline of VM with colocalization of TC cells, axons, and terminals only in VM. Yellow arrows highlight colocalized puncta. Scale bar = 100 μm. **(H)** Summary of normalized GFP and mRuby fluorescence in VM, CM, and MD. n = 5 mice. **(I)** Schematic of CTB-647 injection into mPFC, AAV-Chronos-GFP into SNr, and AAV-ChrimsonR-tdTomato into CbN. **(J)** Representative, CCF-aligned, heatmaps of mPFC, SNr, and CbN injection sites with respective regions of interest outlined in white. Scale bar = 500 μm. Heatmap key shows fluorescence value. **(K)** *Left,* Representative, CCF-aligned, thalamic section, showing GFP+ SNr axons (green) and tdTomato+ CbN axons (red). VM and VAL are outlined in white. Scale bar = 500 μm. *Right,* Confocal images of magnified yellow box outline of VM and VAL. Scale bar = 250 μm. **(L)** Summary of normalized CTB-647, SNr axons, and CbN axons in VM and VAL. n = 3 mice. Values are mean ± SEM. * = p < 0.05.

As a complementary approach, we used anterograde viruses to map axons from SNr to VM thalamus. We injected AAV-FLEX-GFP-2A-Synaptophysin-mRuby in SNr, expressing GFP in axons, and mRuby conjugated to synaptophysin in terminals (Schwarz *et al*., 2015). To isolate inhibitory and excitatory cells in SNr, we injected virus into either Vglut2-Cre or GAD-Cre mice, respectively (Taniguchi *et al*., 2011; Vong *et al*., 2011). In the same animals, we injected CTB-647 into the mPFC to label TC cells in VM thalamus **(Fig. 3E-F & S3C)**. We observed robust axon labeling in GAD-Cre animals (**Fig. 3G**), suggesting that these inputs are likely inhibitory. Across CCF-aligned thalamus, we found GFP+ and mRuby+ expression in VM thalamus, but not in MD or CM (mRuby+ normalized expression for VM = 129.8 ± 37.3 A.U., MD = 34.0 ± 14.7 A.U., CM = 31.9 ± 14.5 A.U.; n = 5 mice) **(Fig. 3H & S3D)**. In contrast, we observed very few axons in the Vglut2-Cre animals (**Fig. 3SE-F**), indicating a lack of glutamatergic cells in SNr.

To assess other subcortical inputs, we injected CTB-647 into the mPFC and AAV-FLEX-GFP-2A-Synaptophysin-mRuby into CbN, and in this case observed minimal labeling of inputs in ventral VM, and little overlap with mPFC-projecting TC cells (**Fig. S3G-H**). To further examine region-specific targeting of VM, in separate experiments, we injected CTB-647 in mPFC and AAV-Chronos-GFP and AAV-Chrimson-tdTomato in SNr and CbN, respectively (**Fig. 3I-J**). Consistent with our other anatomy, we observe close overlap of SNr axons and mPFC-projecting TC cell labeling in VM. In contrast, we found that CbN inputs are biased to dorsal VM and ventral lateral thalamus (VAL), with minimal overlap with PFC-projecting TC cells (**Fig. 3K**). Within CCF-aligned thalamus, we observed SNr axons in both VM and VAL. SNr GFP+ axons in VM tightly overlapped with retrograde CTB-647+ labeling from mPFC. In contrast, we observed robust tdTomato+ CbN axons in VAL, but very few in VM (CTB-647+ normalized expression for VM = 12.3 ± 2.0 A.U. and VAL = 1.8 ± 0.9 A.U.; GFP+ normalized expression for VM = 15.4 ± 2.6 A.U. and VAL = 14.1 ± 6.7 A.U.; tdTomato+ normalized expression for VM = 3.0 ± 1.7 A.U. and VAL = 23.6 ± 8.2 A.U.; n = 3 mice) (**Fig. 3L)**. Together, these results indicate SNr sends putative inhibitory inputs to VM, which overlap with mPFC-projecting TC cells, suggesting that SNr is a major subcortical input.

### Assessing inhibitory synapses from SNr to mPFC-projecting TC cells in VM

Anatomical tracing identifies possible connections, but cannot characterize the strength, sign, and dynamics of synapses. To better understand how SNr engages TC cells in VM, we next used optogenetics and whole-cell recordings in acute coronal slices (Collins *et al*., 2018). We injected AAV-ChR2-eYFP into the SNr, as well as AAVrg-H2B-tdTomato into the mPFC of wild-type mice **(Fig. 4A)**. After allowing for expression and transport, we then prepared acute coronal slices and made voltage-clamp recordings from tdTomato+ TC cells in VM. To assess the sign and dynamics of inputs, we recorded responses to stimulus trains (five pulses @ 10 Hz with 5 mW of 473 nm light) at the reversal for excitation (E_AMPA_) and in the presence of NMDA receptor blockers (10 µM CPP), which allowed responses mediated by GABAergic inhibition to be isolated. We found SNr inputs evoked robust IPSCs at mPFC-projecting TC cells (IPSC_P1_ = 575.1 ± 247.8 pA; n = 19 cells, 8 mice) **(Fig. 4B)**. These responses were blocked by the GABA-A receptor antagonist Gabazine (10 µM) (94.5 ± 3.4% block; Wilcoxon test: W = -21, p = 0.03; n = 6 cells, 3 mice) **(Fig. 4C)**, indicating that they are primarily mediated by these inhibitory receptors.

**Figure 4:**
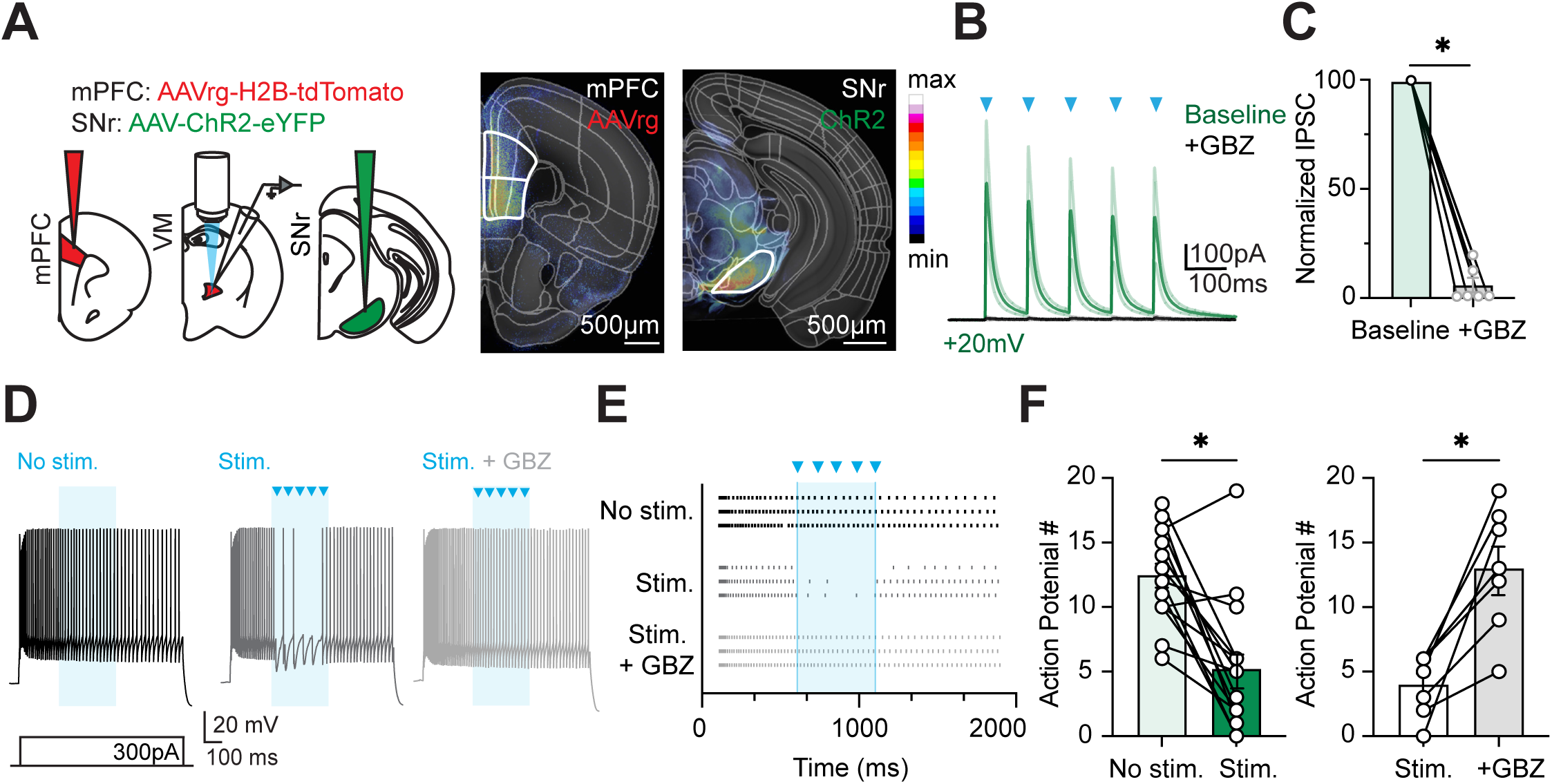
GABAergic SNr inputs inhibit mPFC-projecting TC cells in VM. (**A**) *Left,* Schematic of injections and recording strategy. AAVrg-H2B-tdTomato injected into mPFC and AAV-ChR2-eYFP into SNr of WT mice. Recordings of retrogradely labeled tdTomato+ TC cells in VM. *Middle,* Representative, CCF-aligned, heatmaps of mPFC and *Right,* SNr injection sites with respective regions of interest outlined in white. Scale bar = 500 μm. Heatmap key on right shows fluorescence value. **(B)** SNr-evoked IPSCs at tdTomato+ TC cells in response to 5 x 10 Hz LED stimulation trains (blue triangles) at +20 mV. n = 19 cells, 8 mice. **(C)** Normalized SNr-evoked IPSC with and without Gabazine. 94.5 ± 3.4% block. Wilcoxon test: W = -21, p = 0.03. n = 6 cells, 3 mice. **(D)** Cells are injected with 300 pA of current to evoked steady firing while axons are stimulated with 10 Hz LED stimulation trains (blue triangles). *Left,* No stimulation LED control (black). *Middle,* 5 mW 10 Hz LED stimulation trains pause TC cell firing (dark gray). *Right,* Gabazine blocks inhibition and returns firing to baseline (light gray). **(E)** Raster plots for no stimulation LED control (black), 5 mW 10 Hz LED stimulation (dark gray), and 5 mW 10 Hz LED stimulation + GBZ (light gray). 5 mW 10 Hz LED stimulation trains starting at 600 ms (blue triangles) pause firing. **(F)** *Left,* Summary of number of action potentials fired during 600 – 1100 ms stimulation window for no stimulation LED control and 5 mW stimulation. Wilcoxon test: W = -110, p = 0.0005. n = 15 cells, 8 mice. *Right,* Summary of the number of action potentials during 5 mW LED stimulation with and without Gabazine. Wilcoxon test: W = 28, p = 0.02. n = 7 cells, 7 mice. Values are mean ± SEM. * = p < 0.05.

Having established that SNr makes inhibitory connections onto TC cells, we next evaluated if these inputs are sufficient to pause action potential firing. In separate experiments, we performed similar injections of AAV-ChR2-YFP and AAVrg-H2B-tdTomato but now recorded in current-clamp. To assay for any effects of inhibition, we first injected 2,000 ms depolarizing current steps to drive AP firing of TC cells in VM. We interleaved trains of light (5 mW stimulation) and found that SNr inputs are sufficient to completely pause firing (No stim. = 12.5 ± 0.9 APs, 5 mW stim. = 5.2 ± 1.3 APs; Wilcoxon test: W = -110, p = 0.0005; n = 15 cells, 8 mice) **(Fig. 4D-F).** This strong suppression was also blocked by 10 µM Gabazine (5 mW = 4.0 ± 0.9 APs, 5 mW + Gabazine = 13.0 ± 1.8 APs; Wilcoxon test: W = 28, p = 0.02; n = 7 cells, 7 mice) **(Fig. 4F)**. Together, these results indicate that SNr input to VM is sufficiently strong to pause the activity of mPFC-projecting TC cells, suggesting it can regulate output to the mPFC and ultimately shape cortical dynamics.

### Cortical output of VM TC Cells

Having examined the influence of SNr on TC neurons, we next examined their axonal output to the mPFC. VM is often considered “matrix” thalamus (Jones, 1998; 2001), with axons that arborize broadly across superficial layers of cortex (Harris & Shepherd, 2015). To begin to define the extent of these projections, we injected retrogradely transported AAVrg-Cre in the mPFC, as well as a Cre-dependent anterograde AAV-DIO-GFP into VM (**Fig. 5A**). We observed GFP+ cells in VM, as well as GFP+ axon projections in layer 1 of mPFC, but minimal labeling in the anterolateral motor cortex (ALM) and insular cortex (aIC) (mPFC = 6.3 ± 2.8 A.U., ALM = 0.4 ± 0.2 A.U., aIC = 0.4 ± 0.4 A.U; Friedman test: Q = 6.5, p = 0.04; Dunn’s multiple comparisons test: p = 0.04 for mPFC vs. aIC; n = 4 mice) (**Fig. 5B & 5C**). These findings indicate that mPFC-projecting TC cells in VM project to mPFC, but not more distant parts of the frontal cortex.

**Figure 5:**
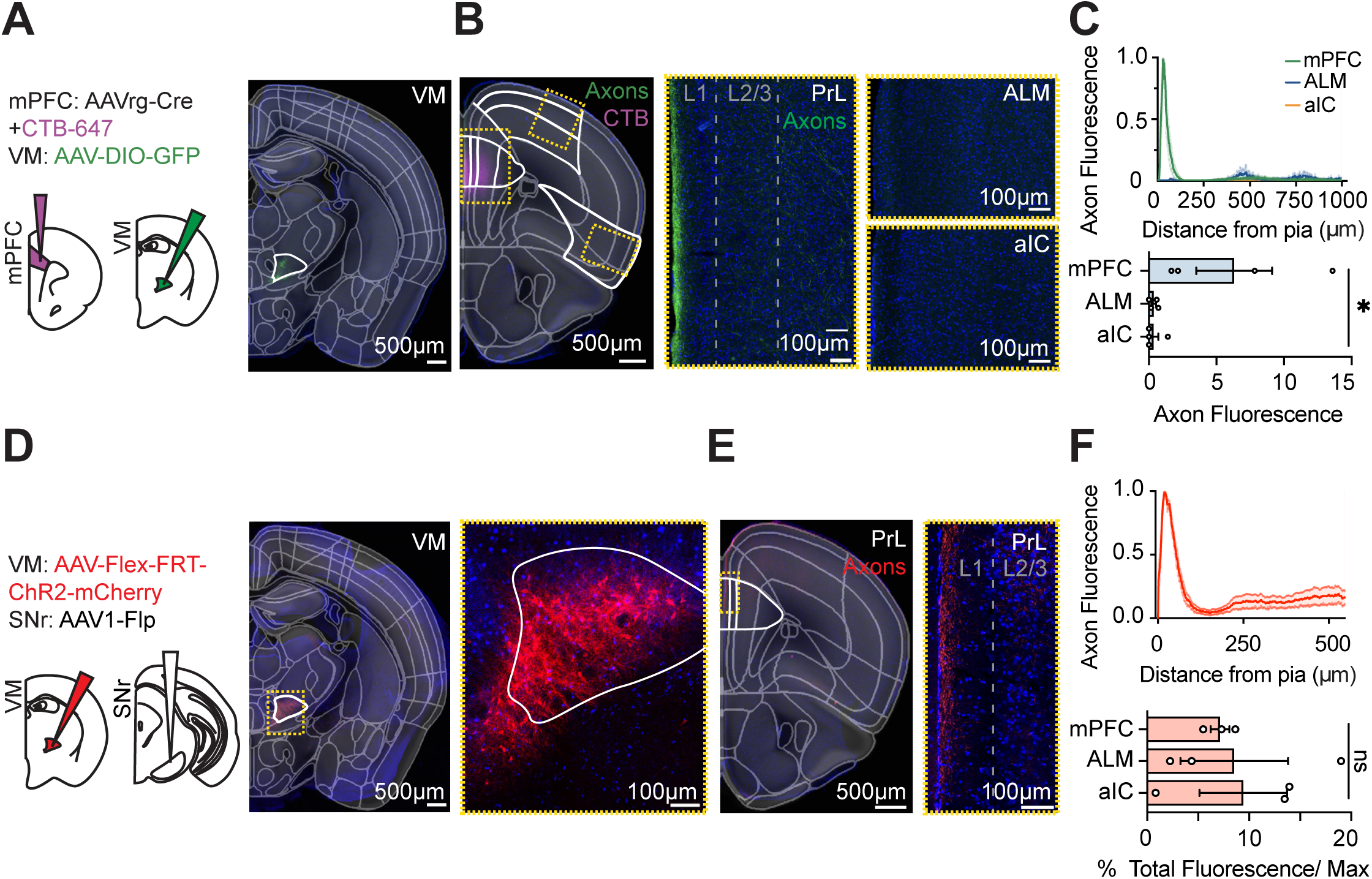
Axonal projections of TC cells in VM to the mPFC. (**A**) *Left,* Schematic of AAVrg-Cre (co-injected with a small volume of CTB-647 to mark injection location) injection into mPFC and AAV-DIO-GFP injection into VM. *Right,* Representative, CCF-aligned, images of VM, showing GFP+ TC cells. VM is outlined in white. Scale bar = 500 μm. **(B)** *Left,* Representative, CCF aligned, image of a cortical coronal section, showing mPFC, ALM, and aIC outlined in white. Scale bar = 500 μm. *Right,* Confocal images of magnified yellow box outlines of mPFC, ALM, and aIC, showing GFP+ axons confined to L1 of mPFC and not present in ALM or aIC. Scale bar = 100 μm. **(C)** *Top,* Summary of axon fluorescence across layers of mPFC, ALM, and aIC. n = 4 mice. *Bottom,* Summary of normalized L1 fluorescence in mPFC, ALM, and aIC. Friedman test: Q = 6.5, p = 0.04; Dunn’s multiple comparisons test: p = 0.04 for mPFC vs. aIC. n = 4 mice. **(D)** *Left,* Schematic of trans-neuronal viral tracing paradigm, with AAV1-Flp and AAV-Flex-FRT-ChR2-mCherry injected into SNr and VM, respectively. *Middle,* Representative, CCF aligned, image of a thalamic coronal section, with VM outlined in white, and mCherry shown in red. Scale bar = 500 μm. *Right,* Confocal image of yellow box outline of VM, where only SNr-contacted VM neurons express mCherry. **(E)** *Left,* Representative, CCF aligned, cortical coronal section, with PrL outlined in white. Scale bar = 500 μm. *Right,* Confocal images of magnified yellow box outline, with mCherry+ axons confined to L1 of PrL. Scale bar = 100 μm. **(F)** *Top,* Summary of axons across layers of mPFC. n = 3 mice. *Bottom,* Summary of normalized bulk L1 fluorescence in mPFC, ALM, and aIC. Friedman test: Q = 0, p > 0.9. n = 3 mice Values are mean ± SEM. * = p < 0.05.

To further assess whether can SNr engage these VM projections to the cortex, we next injected AAV1-Flp into the SNr, which also moves anterogradely and causes expression of Flp in postsynaptic TC cells in VM (**Fig. 5D**). At the same time, we injected Flp-dependent anterograde fluorophore (AAV-Flex-FRT-ChR2-mCherry) into VM, which expresses fluorophore in SNr-contacted TC cells in VM, as well as axons in target brain regions (**Fig. 5D**). With this strategy, we observed prominent axon fluorescence in the prelimbic mPFC (**Fig. 5E**), which was again confined to superficial layer 1 (mPFC = 7.1 ± 0.9 A.U., ALM = 8.5 ± 5.3 A.U., aIC = 9.4 ± 4.3 A.U.; Friedman test: Q = 0, p > 0.9; n = 3 mice) **(Fig. 5F).** In this case, we also observed axon labeling in ALM and aIC, consistent with SNr innervating cells that also project to these parts of the frontal cortex. These anatomical findings confirm that mPFC-projecting TC cells in VM primarily innervate the most superficial layers of the mPFC, suggesting that SNr inputs can engage this pathway.

### SNr input routed through VM contacts engages L1 interneurons in mPFC

We previously found that inhibitory cells expressing Neuron Derived Neurotrophic Factor (NDNF+) are abundant in superficial L1 (Rudy *et al*., 2011; Schuman *et al*., 2019; Anastasiades *et al*., 2021). To study the morphology and physiology of these cells, we injected AAV-FLEX-tdTomato into NDNF-Cre mice (Tasic *et al*., 2016). After allowing for expression, we observed red, tdTomato+ interneurons largely confined to outer L1 **(Fig. 6A)**. In current-clamp recordings, we found cells with local axonal arborizations in L1 **(Fig. 6B)**, which exhibited tendency towards late spiking and high input resistance (R_in_ = 208.4 ± 15.4 MΩ; n = 13 cells, 5 mice) **(Fig. 6C)**. Having defined the properties of these cells, we examined their thalamic innervation by injecting AAV-DIO-eYFP into the mPFC and AAV-ChR2-mCherry into VM **(Fig. 6D).** We then performed voltage-clamp recordings at the reversal for inhibition (E_GABA_), with trains of VM inputs (five pulses @ 10 Hz and 10 mW of 473 nm light), which evoked prominent EPSCs (EPSC_P1_ = 132.2 pA ± 58.0 pA; n = 12 cells, 4 mice) **(Fig. 6E & 6F)**. Lastly, we used trans-neuronal labeling to specifically assess whether SNr inputs are routed via VM thalamus to the mPFC. Using NDNF-Cre mice, we injected AAV1-Flp into the SNr to express Flp in postsynaptic TC cells in VM. In the same animals, we also injected AAV-Flex-FRT-ChR2-mCherry into the VM to express Flp-dependent ChR2 tagged with mCherry. Concurrently, we injected AAV-DIO-eYFP into the mPFC to express Cre-dependent GFP in NDNF+ interneurons **(Fig. 6G)**. In voltage clamp recordings, we found that trains of VM inputs again evoked EPSCs at NDNF+ cells (EPSC_P1_ = 48.9 ± 9.3 pA; n = 11 cells, 4 mice) **(Fig. 6H & 6I**). Together, these data indicate that SNr inputs signal via VM to NDNF+ interneurons in mPFC, forming a polysynaptic circuit that links subcortex through higher-order thalamus to frontal cortex.

**Figure 6:**
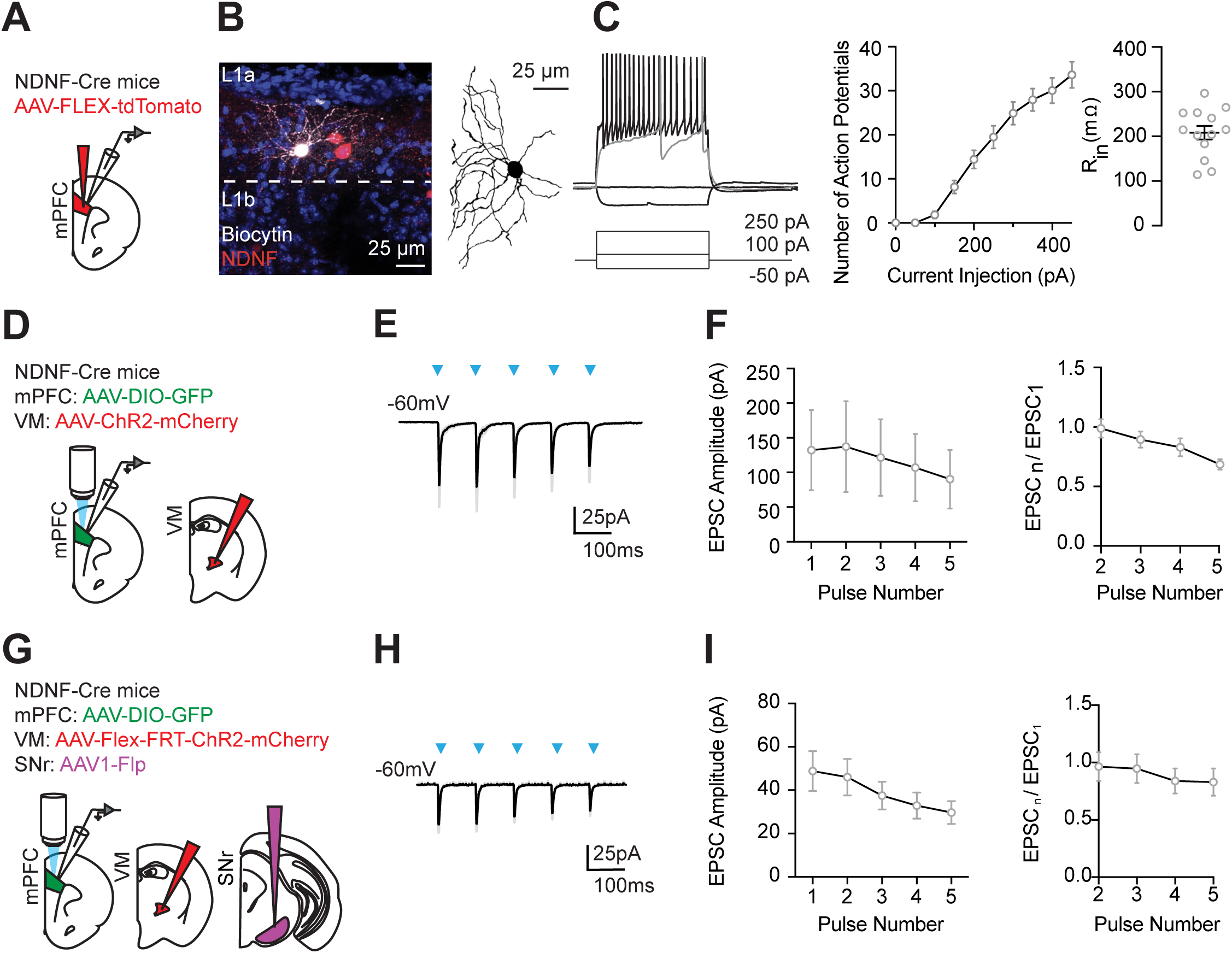
SNr signals via VM to mPFC NDNF+ interneurons in mPFC. (**A**) Schematic of injection paradigm and recording strategy. AAV-FLEX-tdTomato injection into mPFC of NDNF-Cre mice. Recordings from tdTomato+ NDNF+ cells. **(B)** *Left,* Representative image of biocytin-filled (white) tdTomato+ (red) NDNF+ cell in L1a of the mPFC. *Right,* Reconstruction of a tdTomato+ NDNF+ cell in L1a. **(C)** *Left,* Action potential firing of NDNF+ cell in response to current injection of 100 pA (gray trace), 250 pA, and -50 pA (black traces) from -70 mV. *Middle,* Summary input-output curve showing action potential firing in response to increasing injections of current. *Right,* Summary of input resistance. n = 13 cells, 5 mice. **(D)** Schematic of AAV-DIO-GFP injection into mPFC and AAV-ChR2-mCherry into VM of NDNF-Cre mice, and whole-cell recordings of GFP+ cells in the mPFC. **(E)** VM-evoked EPSCs at L1 NDNF+ cells in response to 10 Hz LED stimulation trains (blue triangles) recorded at -60 mV. **(F)** *Left,* Summary of VM-evoked EPSC amplitude versus pulse number at L1 NDNF+ cells. *Right,* Summary of paired-pulse ratio (PPR). n = 12 cells, 4 mice. **(G)** Schematic of trans-neuronal injection paradigm and recording strategy. AAV1-Flp injected into SNr, AAV-Flex-FRT-ChR2-mCherry into VM, and AAV-DIO-GFP into mPFC of NDNF-Cre mice. Recordings from GFP+, NDNF+ cells. **(H)** SNr-contacted VM-evoked EPSCs at GFP+ L1 NDNF+ cells in response to 10 Hz LED stimulation trains (blue triangles) at -60 mV. **(I)** *Left,* Summary of SNr-contacted VM-evoked EPSC amplitude versus pulse number at L1 NDNF+ cells. *Right,* Summary of paired-pulse ratio (PPR). n = 11 cells, 4 mice. Values are mean ± SEM.

## DISCUSSION

We have defined the cellular and synaptic properties of a polysynaptic pathway from the subcortex through VM to the mPFC. We first established that the predominant subcortical input to mPFC-projecting TC cells in VM is the SNr and not CbN or GPi. We then showed that SNr sends robust, GABAergic projections to VM that can strongly inhibit mPFC-projecting TC cells. Finally, we demonstrated how SNr signals can be routed via these TC cells in VM to L1 NDNF+ interneurons in the mPFC. Together, our results illustrate how subcortical brain regions like SNr can mediate activity in VM to engage inhibitory networks in superficial layers of the cortex.

An important feature of our experiments was to define the specific subregion of VM that projects to the mPFC. Our anatomy using anterograde and retrograde labeling confirmed close overlap of mPFC axons and mPFC-projecting TC cells at the ventral border of VM. This recapitulates our previous work showing closed reciprocal loops between VM and mPFC (Collins et al., 2018). We confirmed this retrograde anatomy using three strategies, which showed consistent labeling of TC cells in ventral VM. The clearest labeling was found with rabies virus expressing histone-tagged fluorescent protein, but this approach is challenging to combine with electrophysiology (Callaway, 2008; Callaway & Luo, 2015). CTB labeling closely overlapped with rabies and has two advantages: (1) targeted slice recordings, and (2) anterograde labeling of mPFC axons (Conte *et al*., 2009). AAVrg labeling yielded the sparsest labeling, but still closely overlapped with rabies and RV, and was the best for physiology (Tervo *et al*., 2016; Economo *et al*., 2018).

Previous studies on VM have mostly focused on motor cortex-projecting TC cells in motor thalamus, which are prominent in the dorsal VM and ventrolateral thalamus (VAL) (Sommer, 2003; Hooks *et al*., 2013; Lee *et al*., 2020). In contrast, our results indicate that mPFC-projecting TC cells peak at more anterior locations in the thalamus and are prominent on the ventral VM, and thus substantially offset from motor cortex-projecting TC cells. Once we established the locations of PFC-projecting TC cells in ventral VM, we determined that these cells have radial dendritic arborizations that are similar to those in other thalamic nuclei, including the VAL (Yamamoto *et al*., 1991) and MD (Collins *et al*., 2018; Lyuboslavsky *et al*., 2024), consistent with several previous studies (Peng *et al*., 2021). We also established mPFC-projecting TC cells exhibit burst firing following hyperpolarized potentials, a canonical property of thalamic neurons (Llinas & Jahnsen, 1982; Sherman & Guillery, 1996; Sherman, 2001). These data suggest that distinct TC cells exist within VM, which may send different outputs and may be poised to process different inputs, showing the need for careful labeling in any anatomical and physiological studies.

The unique location of PFC-projecting TC cells in ventral VM suggested they may sample different long-range inputs. Our TRIO anatomy suggested SNr, but not CbN or GPi, is the main subcortical input to mPFC-projecting TC cells in VM. In contrast, previous studies have identified the SNr, CbN, and GPi as the main subcortical inputs to motor thalamus (Sommer, 2003; Lanciego *et al*., 2012; Bosch-Bouju *et al*., 2013; Gao *et al*., 2018; Catanese & Jaeger, 2021). However, work on CbN and GPi inputs to VM typically focus on the dorsal border of VM, proximal to VAL (Gornati *et al*., 2018; Koster & Sherman, 2024). Therefore, lack of substantial CbN and GPi input to mPFC-projecting cells may reflect differences in the dorsal-ventral location of TC cells in VM thalamus.

Reciprocal connections between motor thalamus and cortex have been extensively studied (Oh *et al*., 2014; Shepherd & Yamawaki, 2021; Whyte *et al*., 2024). Notably, reciprocal connections with the mPFC (Collins *et al*., 2018) and motor cortex (Hooks *et al*., 2013; Yamawaki & Shepherd, 2015) underlie persistent activity that enables executive behaviors or movement-related activity, respectively (Wang *et al*., 2021). We observed strong input from PrL, IL, and anterior cingulate (ACC) to mPFC-projecting TC cells, confirming a closed reciprocal loop between mPFC and VM (Collins *et al*., 2018). Surprisingly, we also observed robust inputs from other cortical areas including orbital frontal cortex (OFC), insular cortex (aIC), and motor cortex (MO). Inputs from these cortical areas to mPFC-projecting cells suggests additional “open” loops between cortical different cortical areas and VM (Brown *et al*., 2020), which will be interesting to study in the future.

We used complementary viral strategies and transgenic mice to further characterize anatomical connections from SNr to TC cells in VM. Using Ai14 mice with AAV1-Cre indicated mPFC-projecting, we found SNr-contacted TC cells are confined to ventral VM, as in previous studies (Lee *et al*., 2020). Using Vglut2-Cre and GAD-Cre mice with Cre-dependent viruses indicated GABAergic but not glutamatergic fibers in VM (Liu *et al*., 2020). These data are consistent with SNr being a predominantly GABAergic nucleus with long-range inhibitory output to thalamus (Zhou & Lee, 2011; Catanese & Jaeger, 2021; McElvain *et al*., 2021) In contrast, CbN input does not contact mPFC-projecting VM TC cells, consistent with our other anatomy, but in contrast to previous results in motor thalamus (Proville *et al*., 2014; Gao *et al*., 2018; Gornati *et al*., 2018).

To characterize the sign, dynamics, and receptor contributions of SNr inputs to PFC-projecting TC cells in VM, we used slice physiology. Consistent with prominent GABAergic cells in SNr (Hanaway *et al*., 1970; Richards *et al*., 1997; Zhou & Lee, 2011), we found SNr inputs are inhibitory and depressing, similar to equivalent studies in motor thalamus (Antal *et al*., 2014; Edgerton & Jaeger, 2014; Koster & Sherman, 2024). Consistent with a lack of glutamatergic cells in SNr (Gonzalez-Hernandez & Rodriguez, 2000; McElvain *et al*., 2021), we found no excitatory inputs to TC cells in VM. Ultimately, SNr input is sufficiently strong to pause action potential firing of TC cells, which depends on GABA-A receptors, similar to motor thalamus (Cope *et al*., 2005).

Recent results in motor thalamus indicate that CbN and SNr appear to converge onto TC cells on the dorsal border of VM (Alonso-Martinez *et al*., 2023). These findings suggest distinct subcortical inputs may be processed by individual TC cells before forwarding to motor cortex. However, the lack of prominent CbN input to mPFC-projecting TC cells suggests that comparable processing does not occur in ventral VM. Instead, our anatomy data suggests that the main excitatory input is corticothalamic connections, suggesting that SNr may act as an inhibitory break on cortico - thalamo-cortical loops, potentially pausing persistent activity (Koster & Sherman, 2024).

After characterizing SNr inputs to mPFC-projecting TC cells, we also investigated their axonal projections to the mPFC. VM is defined as “matrix” thalamus, with TC cells sending diffuse projections across the cortex to promote arousal (Jones, 1998; Rubio-Garrido *et al*., 2007). Previous work suggests that a single VM cell could target many cortical regions with its diffuse axon projections (Clasca *et al*., 2012). However, we observed axons from TC cells only in PFC, and not in other cortical regions like ALM and aIC. One explanation is that mPFC-projecting VM cells may function more like “core” thalamus, with more discrete projections. Another possibility is that projections to other cortical regions are averaged out; for example, if some PFC-projecting TC cells project to ALM, and others to aIC, they would appear small when averaged across slices.

Having examined the anatomy of TC output, we used electrophysiology to assess functional connections in the mPFC. We previously showed VM primarily targets NDNF+ interneurons in the most superficial layers of the cortex (Anastasiades *et al*., 2021). We recapitulated these results and observed dense axon coverage of L1 of the mPFC and excitatory input to NDNF+ cells. With transsynaptic optogenetics, we also found that VM neurons contacted by SNr also innervate NDNF+ interneurons, indicating signals are transmitted from the SNr to the mPFC via VM.

Ultimately, we determined that SNr is a potent inhibitory input to TC cells in VM, which themselves activate interneurons in the PFC. Our findings make several testable predictions for future studies: First, we predict that SNr inhibition acts as an inhibitory brake on TC cells in VM, including reciprocal corticothalamic circuits (Catanese & Jaeger, 2021; Koster & Sherman, 2024). Second, we predict that inhibition of VM by SNr leads to reduced drive of NDNF+ cells in the mPFC (Abs *et al*., 2018; Thomas *et al*., 2023; Hartung *et al*., 2024). Third, because NDNF+ cells inhibit other interneurons, we predict that activation of SNr in turn disinhibits the mPFC (Cohen-Kashi Malina *et al*., 2021). Because these circuits are both polysynaptic and long-range, these predictions are challenging to test in slices, but may be addressed in the intact brain in future studies.

In summary, we have characterized how subcortical input from the SNr is transmitted through TC cells in VM to L1 interneurons in the mPFC. We demonstrate SNr is a robust GABAergic input on to mPFC-projecting TC cells and acts through GABA_A_ receptors. Our results contrast with previous findings that model the convergence of several subcortical inputs, including SNr with CbN and GPi in motor thalamus. Lastly, our findings highlight how the subcortical brain regions can act through higher-order thalamus to shape cortical circuits and ultimately influence behavior.

## Acknowledgements

We thank members of the Carter lab and the U19 thalamus team for helpful discussions and comments on the manuscript. This work was supported by NSF 25-A0-00-1015523 (SMC), NIMH R01 MH085974 (AGC), and NINDS U19 NS123714 (AGC). The authors have no financial conflicts of interest.

---

**Supplemental Figure 2-1:**
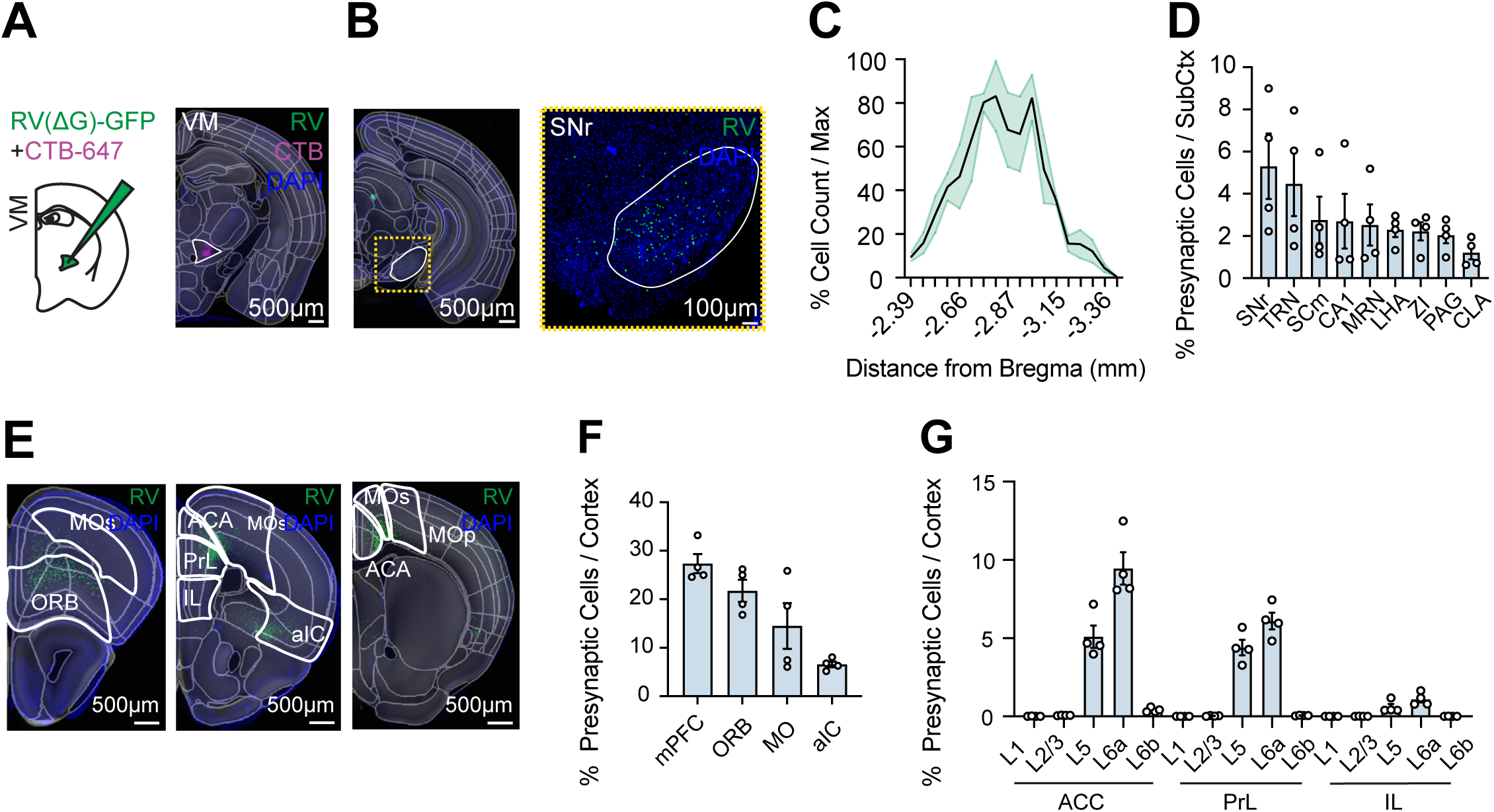
Further characterization of presynaptic cells in TRIO anatomy. **(A)** *Left,* Schematic of RV(ΔG)-GFP injection (co-injected with a small volume of CTB-647 to label injection site) into mPFC. *Right,* Representative, CCF-aligned, thalamic coronal section of VM injection site with GFP+ (green) and CTB-647 (magenta) cells. VM is outlined in white. Scale bar = 500 μm. **(B)** *Left,* Representative, CCF-aligned, SNr coronal section, showing retrogradely labeled GFP+ (green) presynaptic cells. SNr is outlined in white. Scale bar = 500 μm. *Right,* Cropped image of magnified yellow box outline of SNr. Scale bar = 100 μm. **(C)** Distribution of retrogradely labeled, GFP+ cells in SNr across anterior-posterior (A-P) axis, normalized to total cells. A-P coordinate are millimeters from Bregma. n = 4 mice. **(D)** Summary of subcortical inputs as percentage of total cells in subcortical structures. SNr, substantia nigra pars reticulata; TRN, reticular nucleus of the thalamus; SCm, superior colliculus; CA1, Hippocampal field CA1; MRN, median raphe nucleus; LHA, lateral hypothalamic area; ZI, zona incerta; PAG; periaqueductal gray; CLA, claustrum; CLA. n = 4 mice. **(E)** Representative, CCF-aligned, cortical coronal sections, showing GFP+ presynaptic cells in different cortical areas. Scale bar = 500 μm. PrL, prelimbic cortex; IL, infralimbic cortex; ACA, anterior cingulate; ORB, orbital cortex; MOp, primary motor cortex; MOs, secondary motor cortex; aIC, insular cortex. **(F)** Summary of top cortical cells normalized to total presynaptic cells. mPFC, ORB, MO, and aIC. **(G)** Summary of cortical cell count across layers of ACC, PrL, and IL.

**Supplemental Figure 3-1.**
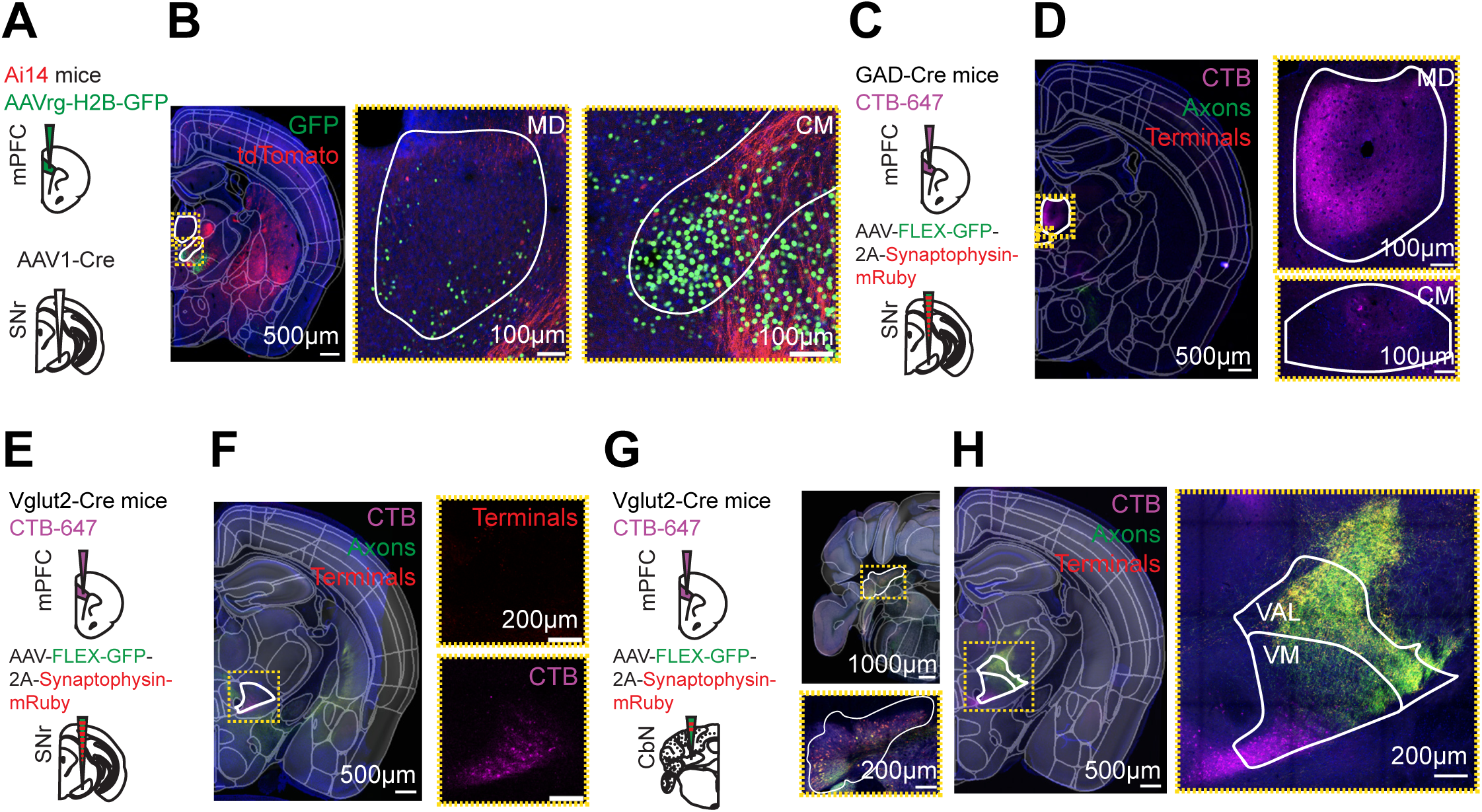
Further characterization of subcortical inputs to VM. **(A)** Schematic of AAVrg-H2B-GFP injection into mPFC and AAV1-Cre injection into SNr of Ai14 mice. **(B)** *Left,* Representative, CCF-aligned, thalamic coronal section, showing retrogradely labeled GFP+ TC cells (green), but few trans-neuronally labeled tdTomato+ cells (red) in MD and CM. Scale bar = 500 μm. *Middle,* Confocal image of magnified yellow box outline of MD and *Right,* CM. Scale bar = 100 μm. **(C)** Schematic of CTB-647 injection into mPFC and AAV-FLEX-GFP-2A-Synaptophysin-mRuby injection into SNr of GAD-Cre mice. **(D)** *Left,* Representative, CCF-aligned, thalamic coronal section, showing retrogradely infected TC cells (magenta), GFP+ SNr axons (green), and mRuby+ SNr terminals (red). MD and CM are outlined in white. Scale bar = 500 μm. *Right,* Confocal image of magnified yellow box outline of MD and CM. Retrogradely infected TC cells (magenta), GFP+ SNr axons (green), and mRuby+ SNr terminals (red). Scale bar = 100 μm. **(E)** Schematic of CTB-647 injection into mPFC and AAV-FLEX-GFP-2A-Synaptophysin-mRuby injection into SNr of a Vglut2-Cre mice. **(F)** *Left,* Representative, CCF-aligned, thalamic coronal section, showing retrogradely infected TC cells (magenta). VM outlined in white. Scale bar = 500 μm. *Right,* Confocal images of magnified yellow box of VM, showing retrogradely infected TC cells, but no axon terminals labeled. Scale bar = 200 μm. **(G)** *Left,* Schematic of CTB-647 injection into mPFC and AAV-FLEX-GFP-2A-Synaptophysin-mRuby injection into CbN of Vglut2-Cre mice. *Right top,* Representative, CCF-aligned, cerebellar coronal section, showing injection site in the CbN. Scale bar = 1000 μm. *Right bottom,* Confocal image of magnified yellow box of CbN. Scale bar = 200 μm. **(H)** *Left,* Representative, CCF aligned, thalamic coronal section, showing TC cells (magenta), GFP+ CbN axons (green), and mRuby+ CbN terminals (red). VM and VAL are outlined in white. Scale bar = 500 μm. *Right,* Confocal image of magnified yellow box outline of VM and VAL. Scale bar = 200 μm.

## REFERENCES

1. Abs, E., Poorthuis, R.B., Apelblat, D., Muhammad, K., Pardi, M.B., Enke, L., Kushinsky, D., Pu, D.L., Eizinger, M.F., Conzelmann, K.K., Spiegel, I. & Letzkus, J.J. (2018) Learning-Related Plasticity in Dendrite-Targeting Layer 1 Interneurons. Neuron, 100, 684–699 e686.

2. Alonso-Martinez, C., Rubio-Teves, M., Porrero, C., Clasca, F. & Casas-Torremocha, D. (2023) Cerebellar and basal ganglia inputs define three main nuclei in the mouse ventral motor thalamus. Front Neuroanat, 17, 1242839.

3. Anastasiades, P.G., Collins, D.P. & Carter, A.G. (2021) Mediodorsal and Ventromedial Thalamus Engage Distinct L1 Circuits in the Prefrontal Cortex. Neuron, 109, 314–330 e314.

4. Antal, M., Beneduce, B.M. & Regehr, W.G. (2014) The substantia nigra conveys target-dependent excitatory and inhibitory outputs from the basal ganglia to the thalamus. J Neurosci, 34, 8032–8042.

5. Aoki, S., Smith, J.B., Li, H., Yan, X., Igarashi, M., Coulon, P., Wickens, J.R., Ruigrok, T.J. & Jin, X. (2019) An open cortico-basal ganglia loop allows limbic control over motor output via the nigrothalamic pathway. Elife, 8.

6. Beckstead, R.M., Domesick, V.B. & Nauta, W.J. (1979) Efferent connections of the substantia nigra and ventral tegmental area in the rat. Brain Res, 175, 191–217.

7. Bosch-Bouju, C., Hyland, B.I. & Parr-Brownlie, L.C. (2013) Motor thalamus integration of cortical, cerebellar and basal ganglia information: implications for normal and parkinsonian conditions. Front Comput Neurosci, 7, 163.

8. Brown, J.W., Taheri, A., Kenyon, R.V., Berger-Wolf, T.Y. & Llano, D.A. (2020) Signal Propagation via Open-Loop Intrathalamic Architectures: A Computational Model. eNeuro, 7.

9. Callaway, E.M. (2008) Transneuronal circuit tracing with neurotropic viruses. Curr Opin Neurobiol, 18, 617–623.

10. Callaway, E.M. & Luo, L. (2015) Monosynaptic Circuit Tracing with Glycoprotein-Deleted Rabies Viruses. J Neurosci, 35, 8979–8985.

11. Catanese, J. & Jaeger, D. (2021) Premotor Ramping of Thalamic Neuronal Activity Is Modulated by Nigral Inputs and Contributes to Control the Timing of Action Release. J Neurosci, 41, 1878–1891.

12. Chalifoux, J.R. & Carter, A.G. (2010) GABAB receptors modulate NMDA receptor calcium signals in dendritic spines. Neuron, 66, 101–113.

13. Chen, T.W., Li, N., Daie, K. & Svoboda, K. (2017) A Map of Anticipatory Activity in Mouse Motor Cortex. Neuron, 94, 866–879 e864.

14. Chen, Z., Wimmer, R.D., Wilson, M.A. & Halassa, M.M. (2015) Thalamic Circuit Mechanisms Link Sensory Processing in Sleep and Attention. Front Neural Circuits, 9, 83.

15. Clasca, F., Rubio-Garrido, P. & Jabaudon, D. (2012) Unveiling the diversity of thalamocortical neuron subtypes. Eur J Neurosci, 35, 1524–1532.

16. Cohen-Kashi Malina, K., Tsivourakis, E., Kushinsky, D., Apelblat, D., Shtiglitz, S., Zohar, E., Sokoletsky, M., Tasaka, G.I., Mizrahi, A., Lampl, I. & Spiegel, I. (2021) NDNF interneurons in layer 1 gain-modulate whole cortical columns according to an animal’s behavioral state. Neuron, 109, 2150–2164 e2155.

17. Collins, D.P., Anastasiades, P.G., Marlin, J.J. & Carter, A.G. (2018) Reciprocal Circuits Linking the Prefrontal Cortex with Dorsal and Ventral Thalamic Nuclei. Neuron, 98, 366–379 e364.

18. Conte, W.L., Kamishina, H. & Reep, R.L. (2009) The efficacy of the fluorescent conjugates of cholera toxin subunit B for multiple retrograde tract tracing in the central nervous system. Brain Struct Funct, 213, 367–373.

19. Cope, D.W., Hughes, S.W. & Crunelli, V. (2005) GABAA receptor-mediated tonic inhibition in thalamic neurons. J Neurosci, 25, 11553–11563.

20. Cruikshank, S.J., Ahmed, O.J., Stevens, T.R., Patrick, S.L., Gonzalez, A.N., Elmaleh, M. & Connors, B.W. (2012) Thalamic control of layer 1 circuits in prefrontal cortex. J Neurosci, 32, 17813–17823.

21. Deniau, J.M., Menetrey, A. & Thierry, A.M. (1994) Indirect nucleus accumbens input to the prefrontal cortex via the substantia nigra pars reticulata: a combined anatomical and electrophysiological study in the rat. Neuroscience, 61, 533–545.

22. Economo, M.N., Viswanathan, S., Tasic, B., Bas, E., Winnubst, J., Menon, V., Graybuck, L.T., Nguyen, T.N., Smith, K.A., Yao, Z., Wang, L., Gerfen, C.R., Chandrashekar, J., Zeng, H., Looger, L.L. & Svoboda, K. (2018) Distinct descending motor cortex pathways and their roles in movement. Nature, 563, 79–84.

23. Edgerton, J.R. & Jaeger, D. (2014) Optogenetic activation of nigral inhibitory inputs to motor thalamus in the mouse reveals classic inhibition with little potential for rebound activation. Front Cell Neurosci, 8, 36.

24. Euston, D.R., Gruber, A.J. & McNaughton, B.L. (2012) The role of medial prefrontal cortex in memory and decision making. Neuron, 76, 1057–1070.

25. Gao, Z., Davis, C., Thomas, A.M., Economo, M.N., Abrego, A.M., Svoboda, K., De Zeeuw, C.I. & Li, N. (2018) A cortico-cerebellar loop for motor planning. Nature, 563, 113–116.

26. Glenn, L.L. & Steriade, M. (1982) Discharge rate and excitability of cortically projecting intralaminar thalamic neurons during waking and sleep states. J Neurosci, 2, 1387–1404.

27. Gong, S., Zheng, C., Doughty, M.L., Losos, K., Didkovsky, N., Schambra, U.B., Nowak, N.J., Joyner, A., Leblanc, G., Hatten, M.E. & Heintz, N. (2003) A gene expression atlas of the central nervous system based on bacterial artificial chromosomes. Nature, 425, 917–925.

28. Gonzalez-Hernandez, T. & Rodriguez, M. (2000) Compartmental organization and chemical profile of dopaminergic and GABAergic neurons in the substantia nigra of the rat. J Comp Neurol, 421, 107–135.

29. Gornati, S.V., Schafer, C.B., Eelkman Rooda, O.H.J., Nigg, A.L., De Zeeuw, C.I. & Hoebeek F.E. (2018) Differentiating Cerebellar Impact on Thalamic Nuclei. Cell Rep, 23, 2690–2704.

30. Guo, K., Yamawaki, N., Svoboda, K. & Shepherd, G.M.G. (2018) Anterolateral Motor Cortex Connects with a Medial Subdivision of Ventromedial Thalamus through Cell Type-Specific Circuits, Forming an Excitatory Thalamo-Cortico-Thalamic Loop via Layer 1 Apical Tuft Dendrites of Layer 5B Pyramidal Tract Type Neurons. J Neurosci, 38, 8787–8797.

31. Guo, Z.V., Inagaki, H.K., Daie, K., Druckmann, S., Gerfen, C.R. & Svoboda, K. (2017) Maintenance of persistent activity in a frontal thalamocortical loop. Nature, 545, 181–186.

32. Halassa, M.M. & Sherman, S.M. (2019) Thalamocortical Circuit Motifs: A General Framework. Neuron, 103, 762–770.

33. Hanaway, J., McConnell, J.A. & Netsky, M.G. (1970) Cytoarchitecture of the substantia nigra in the rat. Am J Anat, 129, 417–437.

34. Harris, K.D. & Shepherd, G.M. (2015) The neocortical circuit: themes and variations. Nat Neurosci, 18, 170–181.

35. Hartung, J., Schroeder, A., Perez Vazquez, R.A., Poorthuis, R.B. & Letzkus, J.J. (2024) Layer 1 NDNF interneurons are specialized top-down master regulators of cortical circuits. Cell Rep, 43, 114212.

36. Honjoh, S., Sasai, S., Schiereck, S.S., Nagai, H., Tononi, G. & Cirelli, C. (2018) Regulation of cortical activity and arousal by the matrix cells of the ventromedial thalamic nucleus. Nat Commun, 9, 2100.

37. Hooks, B.M., Mao, T., Gutnisky, D.A., Yamawaki, N., Svoboda, K. & Shepherd, G.M. (2013) Organization of cortical and thalamic input to pyramidal neurons in mouse motor cortex. J Neurosci, 33, 748–760.

38. Huang, A.S., Rogers, B.P. & Woodward, N.D. (2019) Disrupted modulation of thalamus activation and thalamocortical connectivity during dual task performance in schizophrenia. Schizophr Res, 210, 270–277.

39. Huang, S., Wu, S.J., Sansone, G., Ibrahim, L.A. & Fishell, G. (2024) Layer 1 neocortex: Gating and integrating multidimensional signals. Neuron, 112, 184–200.

40. Ibrahim, L.A., Huang, S., Fernandez-Otero, M., Sherer, M., Qiu, Y., Vemuri, S., Xu, Q., Machold, R., Pouchelon, G., Rudy, B. & Fishell, G. (2021) Bottom-up inputs are required for establishment of top-down connectivity onto cortical layer 1 neurogliaform cells. Neuron, 109, 3473–3485 e3475.

41. Iidaka, T. (2021) Fluctuations in Arousal Correlate with Neural Activity in the Human Thalamus. Cereb Cortex Commun, 2, tgab055.

42. Inagaki, H.K., Chen, S., Ridder, M.C., Sah, P., Li, N., Yang, Z., Hasanbegovic, H., Gao, Z., Gerfen, C.R. & Svoboda, K. (2022) A midbrain-thalamus-cortex circuit reorganizes cortical dynamics to initiate movement. Cell, 185, 1065–1081 e1023.

43. Jones, E.G. (1998) Viewpoint: the core and matrix of thalamic organization. Neuroscience, 85, 331–345.

44. Jones, E.G. (2001) The thalamic matrix and thalamocortical synchrony. Trends Neurosci, 24, 595–601.

45. Kamalova, A., Manoocheri, K., Liu, X., Casello, S.M., Huang, M., Baimel, C., Jang, E.V., Anastasiades, P.G., Collins, D.P. & Carter, A.G. (2024) CCK+ interneurons contribute to thalamus-evoked feed-forward inhibition in prelimbic prefrontal cortex. J Neurosci.

46. Kase, D., Uta, D., Ishihara, H. & Imoto, K. (2015) Inhibitory synaptic transmission from the substantia nigra pars reticulata to the ventral medial thalamus in mice. Neurosci Res, 97, 26–35.

47. Klein-Flugge, M.C., Bongioanni, A. & Rushworth, M.F.S. (2022) Medial and orbital frontal cortex in decision-making and flexible behavior. Neuron, 110, 2743–2770.

48. Komiyama, T., Sato, T.R., O’Connor, D.H., Zhang, Y.X., Huber, D., Hooks, B.M., Gabitto, M. & Svoboda, K. (2010) Learning-related fine-scale specificity imaged in motor cortex circuits of behaving mice. Nature, 464, 1182–1186.

49. Koster, K.P. & Sherman, S.M. (2024) Convergence of inputs from the basal ganglia with layer 5 of motor cortex and cerebellum in mouse motor thalamus. Elife, 13.

50. Kros, L., Eelkman Rooda, O.H., Spanke, J.K., Alva, P., van Dongen, M.N., Karapatis, A., Tolner, E.A., Strydis, C., Davey, N., Winkelman, B.H., Negrello, M., Serdijn, W.A., Steuber, V., van den Maagdenberg, A.M., De Zeeuw, C.I. & Hoebeek, F.E. (2015) Cerebellar output controls generalized spike-and-wave discharge occurrence. Ann Neurol, 77, 1027–1049.

51. Kuramoto, E., Ohno, S., Furuta, T., Unzai, T., Tanaka, Y.R., Hioki, H. & Kaneko, T. (2015) Ventral medial nucleus neurons send thalamocortical afferents more widely and more preferentially to layer 1 than neurons of the ventral anterior-ventral lateral nuclear complex in the rat. Cereb Cortex, 25, 221–235.

52. Kuramoto, E., Pan, S., Furuta, T., Tanaka, Y.R., Iwai, H., Yamanaka, A., Ohno, S., Kaneko, T., Goto, T. & Hioki, H. (2017) Individual mediodorsal thalamic neurons project to multiple areas of the rat prefrontal cortex: A single neuron-tracing study using virus vectors. J Comp Neurol, 525, 166–185.

53. Lanciego, J.L., Luquin, N. & Obeso, J.A. (2012) Functional neuroanatomy of the basal ganglia. Cold Spring Harb Perspect Med, 2, a009621.

54. Lee, J., Wang, W. & Sabatini, B.L. (2020) Anatomically segregated basal ganglia pathways allow parallel behavioral modulation. Nat Neurosci, 23, 1388–1398.

55. Liu, D., Li, W., Ma, C., Zheng, W., Yao, Y., Tso, C.F., Zhong, P., Chen, X., Song, J.H., Choi, W., Paik, S.B., Han, H. & Dan, Y. (2020) A common hub for sleep and motor control in the substantia nigra. Science, 367, 440–445.

56. Llinas, R. & Jahnsen, H. (1982) Electrophysiology of mammalian thalamic neurones in vitro. Nature, 297, 406–408.

57. Lobb, C.J. & Jaeger, D. (2015) Bursting activity of substantia nigra pars reticulata neurons in mouse parkinsonism in awake and anesthetized states. Neurobiol Dis, 75, 177–185.

58. Lyuboslavsky, P., Ordemann, G.J., Kizimenko, A. & Brumback, A.C. (2024) Two contrasting mediodorsal thalamic circuits target the mouse medial prefrontal cortex. J Neurophysiol, 131, 876–890.

59. Madisen, L., Zwingman, T.A., Sunkin, S.M., Oh, S.W., Zariwala, H.A., Gu, H., Ng, L.L., Palmiter, R.D., Hawrylycz, M.J., Jones, A.R., Lein, E.S. & Zeng, H. (2010) A robust and high-throughput Cre reporting and characterization system for the whole mouse brain. Nat Neurosci, 13, 133–140.

60. McElvain, L.E., Chen, Y., Moore, J.D., Brigidi, G.S., Bloodgood, B.L., Lim, B.K., Costa, R.M. & Kleinfeld, D. (2021) Specific populations of basal ganglia output neurons target distinct brain stem areas while collateralizing throughout the diencephalon. Neuron, 109, 1721–1738 e1724.

61. Oh, S.W., Harris, J.A., Ng, L., Winslow, B., Cain, N., Mihalas, S., Wang, Q., Lau, C., Kuan, L., Henry, A.M., Mortrud, M.T., Ouellette, B., Nguyen, T.N., Sorensen, S.A., Slaughterbeck, C.R., Wakeman, W., Li, Y., Feng, D., Ho, A., Nicholas, E., Hirokawa, K.E., Bohn, P., Joines, K.M., Peng, H., Hawrylycz, M.J., Phillips, J.W., Hohmann, J.G., Wohnoutka, P., Gerfen, C.R., Koch, C., Bernard, A., Dang, C., Jones, A.R. & Zeng, H. (2014) A mesoscale connectome of the mouse brain. Nature, 508, 207–214.

62. Peng, H., Xie, P., Liu, L., Kuang, X., Wang, Y., Qu, L., Gong, H., Jiang, S., Li, A., Ruan, Z., Ding, L., Yao, Z., Chen, C., Chen, M., Daigle, T.L., Dalley, R., Ding, Z., Duan, Y., Feiner, A., He, P., Hill, C., Hirokawa, K.E., Hong, G., Huang, L., Kebede, S., Kuo, H.C., Larsen, R., Lesnar, P., Li, L., Li, Q., Li, X., Li, Y., Li, Y., Liu, A., Lu, D., Mok, S., Ng, L., Nguyen, T.N., Ouyang, Q., Pan, J., Shen, E., Song, Y., Sunkin, S.M., Tasic, B., Veldman, M.B., Wakeman, W., Wan, W., Wang, P., Wang, Q., Wang, T., Wang, Y., Xiong, F., Xiong, W., Xu, W., Ye, M., Yin, L., Yu, Y., Yuan, J., Yuan, J., Yun, Z., Zeng, S., Zhang, S., Zhao, S., Zhao, Z., Zhou, Z., Huang, Z.J., Esposito, L., Hawrylycz, M.J., Sorensen, S.A., Yang, X.W., Zheng, Y., Gu, Z., Xie, W., Koch, C., Luo, Q., Harris, J.A., Wang, Y. & Zeng, H. (2021) Morphological diversity of single neurons in molecularly defined cell types. Nature, 598, 174–181.

63. Pologruto, T.A., Sabatini, B.L. & Svoboda, K. (2003) ScanImage: flexible software for operating laser scanning microscopes. Biomed Eng Online, 2, 13.

64. Proville, R.D., Spolidoro, M., Guyon, N., Dugue, G.P., Selimi, F., Isope, P., Popa, D. & Lena, C. (2014) Cerebellum involvement in cortical sensorimotor circuits for the control of voluntary movements. Nat Neurosci, 17, 1233–1239.

65. Richards, C.D., Shiroyama, T. & Kitai, S.T. (1997) Electrophysiological and immunocytochemical characterization of GABA and dopamine neurons in the substantia nigra of the rat. Neuroscience, 80, 545–557.

66. Rubio-Garrido, P., Perez-de-Manzo, F. & Clasca, F. (2007) Calcium-binding proteins as markers of layer-I projecting vs. deep layer-projecting thalamocortical neurons: a double-labeling analysis in the rat. Neuroscience, 149, 242–250.

67. Rudy, B., Fishell, G., Lee, S. & Hjerling-Leffler, J. (2011) Three groups of interneurons account for nearly 100% of neocortical GABAergic neurons. Dev Neurobiol, 71, 45–61.

68. Schuman, B., Machold, R.P., Hashikawa, Y., Fuzik, J., Fishell, G.J. & Rudy, B. (2019) Four Unique Interneuron Populations Reside in Neocortical Layer 1. J Neurosci, 39, 125–139.

69. Schwarz, L.A., Miyamichi, K., Gao, X.J., Beier, K.T., Weissbourd, B., DeLoach, K.E., Ren, J., Ibanes, S., Malenka, R.C., Kremer, E.J. & Luo, L. (2015) Viral-genetic tracing of the input-output organization of a central noradrenaline circuit. Nature, 524, 88–92.

70. Shepherd, G.M.G. & Yamawaki, N. (2021) Untangling the cortico-thalamo-cortical loop: cellular pieces of a knotty circuit puzzle. Nat Rev Neurosci, 22, 389–406.

71. Sherman, S.M. (2001) Tonic and burst firing: dual modes of thalamocortical relay. Trends Neurosci, 24, 122–126.

72. Sherman, S.M. & Guillery, R.W. (1996) Functional organization of thalamocortical relays. J Neurophysiol, 76, 1367–1395.

73. Sieveritz, B., Garcia-Munoz, M. & Arbuthnott, G.W. (2019) Thalamic afferents to prefrontal cortices from ventral motor nuclei in decision-making. Eur J Neurosci, 49, 646–657.

74. Sommer, M.A. (2003) The role of the thalamus in motor control. Curr Opin Neurobiol, 13, 663–670.

75. Takahashi, N., Moberg, S., Zolnik, T.A., Catanese, J., Sachdev, R.N.S., Larkum, M.E. & Jaeger, D. (2021) Thalamic input to motor cortex facilitates goal-directed action initiation. Curr Biol, 31, 4148–4155 e4144.

76. Taniguchi, H., He, M., Wu, P., Kim, S., Paik, R., Sugino, K., Kvitsiani, D., Fu, Y., Lu, J., Lin, Y., Miyoshi, G., Shima, Y., Fishell, G., Nelson, S.B. & Huang, Z.J. (2011) A resource of Cre driver lines for genetic targeting of GABAergic neurons in cerebral cortex. Neuron, 71, 995–1013.

77. Tasic, B., Menon, V., Nguyen, T.N., Kim, T.K., Jarsky, T., Yao, Z., Levi, B., Gray, L.T., Sorensen, S.A., Dolbeare, T., Bertagnolli, D., Goldy, J., Shapovalova, N., Parry, S., Lee, C., Smith, K., Bernard, A., Madisen, L., Sunkin, S.M., Hawrylycz, M., Koch, C. & Zeng, H. (2016) Adult mouse cortical cell taxonomy revealed by single cell transcriptomics. Nat Neurosci, 19, 335–346.

78. Tervo, D.G., Hwang, B.Y., Viswanathan, S., Gaj, T., Lavzin, M., Ritola, K.D., Lindo, S., Michael, S., Kuleshova, E., Ojala, D., Huang, C.C., Gerfen, C.R., Schiller, J., Dudman, J.T., Hantman, A.W., Looger, L.L., Schaffer, D.V. & Karpova, A.Y. (2016) A Designer AAV Variant Permits Efficient Retrograde Access to Projection Neurons. Neuron, 92, 372–382.

79. Thomas, A., Yang, W., Wang, C., Tipparaju, S.L., Chen, G., Sullivan, B., Swiekatowski, K., Tatam, M., Gerfen, C. & Li, N. (2023) Superior colliculus bidirectionally modulates choice activity in frontal cortex. Nat Commun, 14, 7358.

80. Varga, C., Sik, A., Lavallee, P. & Deschenes, M. (2002) Dendroarchitecture of relay cells in thalamic barreloids: a substrate for cross-whisker modulation. J Neurosci, 22, 6186–6194.

81. Vong, L., Ye, C., Yang, Z., Choi, B., Chua, S. & Lowell, B.B. (2011) Leptin action on GABAergic neurons prevents obesity and reduces inhibitory tone to POMC neurons. Neuron, 71, 142–154.

82. Wang, Q., Ding, S.L., Li, Y., Royall, J., Feng, D., Lesnar, P., Graddis, N., Naeemi, M., Facer, B., Ho, A., Dolbeare, T., Blanchard, B., Dee, N., Wakeman, W., Hirokawa, K.E., Szafer, A., Sunkin, S.M., Oh, S.W., Bernard, A., Phillips, J.W., Hawrylycz, M., Koch, C., Zeng, H., Harris, J.A. & Ng, L. (2020a) The Allen Mouse Brain Common Coordinate Framework: A 3D Reference Atlas. Cell, 181, 936–953 e920.

83. Wang, Q., Ding, S.L., Li, Y., Royall, J., Feng, D., Lesnar, P., Graddis, N., Naeemi, M., Facer, B., Ho, A., Dolbeare, T., Blanchard, B., Dee, N., Wakeman, W., Hirokawa, K.E., Szafer, A., Sunkin, S.M., Oh, S.W., Bernard, A., Phillips, J.W., Hawrylycz, M., Koch, C., Zeng, H., Harris, J.A. & Ng, L. (2020b) The Allen Mouse Brain Common Coordinate Framework: A 3D Reference Atlas. Cell, 181, 936–953.e920.

84. Wang, Y., Yin, X., Zhang, Z., Li, J., Zhao, W. & Guo, Z.V. (2021) A cortico-basal ganglia-thalamo-cortical channel underlying short-term memory. Neuron, 109, 3486–3499 e3487.

85. Whyte, C.J., Redinbaugh, M.J., Shine, J.M. & Saalmann, Y.B. (2024) Thalamic contributions to the state and contents of consciousness. Neuron, 112, 1611–1625.

86. Xu, P., Chen, A., Li, Y., Xing, X. & Lu, H. (2019) Medial prefrontal cortex in neurological diseases. Physiol Genomics, 51, 432–442.

87. Yamamoto, T., Kishimoto, Y., Yoshikawa, H. & Oka, H. (1991) Intracellular recordings from rat thalamic VL neurons: a study combined with intracellular staining. Exp Brain Res, 87, 245–253.

88. Yamawaki, N. & Shepherd, G.M. (2015) Synaptic circuit organization of motor corticothalamic neurons. J Neurosci, 35, 2293–2307.

89. Zhou, F.M. & Lee, C.R. (2011) Intrinsic and integrative properties of substantia nigra pars reticulata neurons. Neuroscience, 198, 69–94.

90. Zingg, B., Chou, X.L., Zhang, Z.G., Mesik, L., Liang, F., Tao, H.W. & Zhang, L.I. (2017) AAV-Mediated Anterograde Transsynaptic Tagging: Mapping Corticocollicular Input-Defined Neural Pathways for Defense Behaviors. Neuron, 93, 33–47.

